# Type 17 Follicular Helper T (Tfh17) Cells are Superior for Memory Maintenance

**DOI:** 10.1101/2022.07.31.502219

**Authors:** Xin Gao, Kaiming Luo, Diya Wang, Yunbo Wei, Yin Yao, Jun Deng, Yang Yang, Qunxiong Zeng, Xiaoru Dong, Le Xiong, Dongcheng Gong, Lin Lin, Kai Pohl, Shaoling Liu, Yu Liu, Lu Liu, Thi H. O. Nguyen, Lilith F. Allen, Katherine Kedzierska, Yanliang Jin, Meirong Du, Wanping Chen, Liangjing Lu, Nan Shen, Zheng Liu, Ian A. Cockburn, Wenjing Luo, Di Yu

## Abstract

A defining feature of successful vaccination is the ability to induce long-lived antigen-specific memory cells. Follicular helper T (Tfh) cells specialize in providing help to B cells in mounting protective humoral immunity in infection and after vaccination. Memory Tfh cells that retain the CXCR5 expression can confer protection through enhancing humoral response upon antigen re-exposure but how they are maintained is poorly understood. CXCR5^+^ memory Tfh cells in human blood are divided into Tfh1, Tfh2 and Tfh17 cells by the expression of chemokine receptors CXCR3 and CCR6 associated with Th1 and Th17 respectively. Here, we developed a new method to induce Tfh1, Tfh2 and Tfh17-like (iTfh1, iTfh2 and iTfh17) cells *in vitro*. Although all three iTfh subsets efficiently support antibody responses in recipient mice with immediate immunization, iTfh17 cells are superior to iTfh1 and iTfh2 cells in supporting antibody response to a later immunization after extended resting *in vivo* to mimic memory maintenance. Notably, the counterpart human Tfh17 cells are selectively enriched in CCR7^+^ central memory Tfh (Tfh_CM_) with survival and proliferative advantages. Furthermore, the analysis of multiple human cohorts that received different vaccines for HBV, influenza virus, tetanus toxin or measles revealed that vaccine-specific Tfh17 cells outcompete Tfh1 or Tfh2 cells for the persistence in memory phase. Therefore, the complementary mouse and human results showing the advantage of Tfh17 cells in maintenance and memory function supports the notion that Tfh17-induced immunization might be preferable in vaccine development to confer long-term protection.

## Introduction

Follicular helper T (Tfh) cells are the specialized CD4^+^ T cell subset that localize within B cell follicle to assist germinal center (GC) formation, plasma cell differentiation and high-affinity antibody production (1, 2). Identified in the circulation and lymphoid organs post immune response (memory phase), CXCR5^+^ memory Tfh cells rapidly differentiate into mature effector Tfh cells and accelerate antibody response upon antigen re-exposure (3–8).

CXCR5-expressing CD4^+^ T cells circulating in human blood (cTfh) provide important subjects to investigate memory Tfh cells since they have egressed from the site of the immune response at secondary lymphoid organs and can differentiate into effector Tfh cells upon antigen re-exposure (9). Noticeably, cTfh cells are heterogenous and are often classified into subsets by distinct functional markers. For example, cTfh cells are classified into Tfh1 (CXCR3^+^CCR6^-^), Tfh2 (CXCR3^-^CCR6^-^) and Tfh17 (CXCR3^-^CCR6^+^) subsets based on the expression of chemokine receptors CXCR3 and CCR6 associated with Th1 and Th17 respectively. Tfh2 and Tfh17 cells were reported to demonstrate better B cell helper function than Tfh1 cells in culture (10). The increases in Tfh2 and Tfh17 frequencies in autoimmune diseases often correlated with excessive production of pathogenic autoantibodies (11, 12). On the other hand, infections such as HIV and malaria mainly induce the generation of Tfh1 cells (13, 14). In influenza vaccination, it is also the Tfh1 subset that correlates with the titers of protective antibodies (15).

Besides the classification of cTfh into Tfh1, Tfh2 and Tfh17 subsets by the features of lineage polarization, cTfh cells are composed of CCR7^high^PD-1^low^ “central memory (CM)-like” (Tfh_CM_) and CCR7^low^PD-1^high^ “effector memory (EM)-like” (Tfh_EM_) subsets (4), the latter also containing a more active ICOS^+^ population (15–17). CCR7^high^ PD-1^low^ Tfh_CM_ cells are dominant in human blood cTfh cells whereas circulating CCR7^low^PD-1^high^ Tfh_EM_ cells are temporarily induced in immune response and generated from the precursor stage of Tfh differentiation at secondary lymphoid organs (4).

There is little knowledge on the difference of memory Tfh subsets either Tfh1, Tfh2 and Tfh17 or Tfh_EM_ and Tfh_CM_ in memory responses. In this study, we developed a method to induce antigen-specific Tfh1, Tfh2 and Tfh17-like (iTfh1, iTfh2 and iTfh17) mouse cells *in vitro.* iTfh1, iTfh2 and iTfh17 cells showed comparable B-helper function after the adoptive transfer into recipient mice followed by immediate immunization. In contrast, if transferred cells experienced an extended period of resting before re-immunization, iTfh17 cells were superior to iTfh1 and iTfh2 cells in sustaining antibody responses. Human Tfh17 cells represented ∼20% Tfh_EM_ cells but accounted for >50% Tfh_CM_ cells which transcriptionally and phenotypically resemble central memory CD4^+^ T (T_CM_) with better survival and proliferative potential than effector memory (T_EM_) cells (18). In vaccine responses to hepatitis B virus (HBV), influenza, tetanus toxin or measles, the Tfh17 subset in vaccine-specific cTfh cells is preferentially maintained into memory phase and long-lived. Complementary results from mouse and human studies thus suggest Tfh17 cells are superior for memory maintenance, the ability to persist and to support humoral response upon antigen restimulation.

## Results

### The *in vitro* differentiation of induced Tfh1, Tfh2 and Tfh17-like (iTfh1, iTfh2, iTfh17) cells

The classification of Tfh1/2/17 cells based on the expression of CXCR3 and CCR6 markers was established by characterizing human blood memory Tfh cells (10). Such memory Tfh1/2/17 subsets in CD44^+^CXCR5^+^ Tfh cells are low in numbers (**Figure 1A**), thus limiting functional characterization. We modified an established method that induces antigen-specific naïve CD4^+^ T cells, such as OT-II T cells with transgenic TCR specific to ovalbumin (OVA), to differentiate into Tfh cells (iTfh) *in vitro* (19) and induced the individual differentiation into Tfh1/2/17 (iTfh1/2/17) *in vitro*. In addition to IL-6 and IL-21 in the iTfh differentiation method (19), Th1 (IL-12, anti-TGF-β, anti-IL-4), Th17 (TGF-β, anti-IFN-γ, anti-IL-4) and Th2 (IL-4, anti-IFN-γ, anti-TGF-β) polarization cultures were adopted for iTfh1/2/17 induction with lower IL-12, TGF-β or IL-4 concentrations for iTfh1/2/17 induction than those used for canonical iTh1, iTh17 or iTh2 induction (details in the method) (20–22) (**Figure 1B**). iTfh1/2/17 cells expressed higher Tfh-defining markers CXCR5, PD-1 and BCL6 than iTh0/1/2/17 cells (**Figure 1C, D**). The BCL6 expression in iTfh1/2/17 cells was lower than CD44^+^CXCR5^high^PD-1^high^ GC-Tfh cells in immunized mice **(Figure S1A, B)**. Tfh differentiation undergoes a step-by-step process, showing the generation of precursor Tfh cells expressing intermediate levels of BCL6 and subsequent maturation of GC-Tfh cells with the highest BCL6 expression (1, 2). Memory cTfh cells largely originate from precursor Tfh cells and express low levels of BCL6 (4). Given that iTfh1/2/17 cells expressed BCL6 lower than that in GC-Tfh cells and resemble precursor Tfh cells, iTfh1/2/17 cells are suitable to study the function of memory cTfh cells. Importantly, iTfh1/2/17 cells differentially expressed transcription factors T-bet, GATA3 and RORγt (**Figure 1E, F**) and chemokine receptors CXCR3 and CCR6, as their counterpart human Tfh12/17 cells (10) (**Figure 1G, H**).

**Figure 1.**
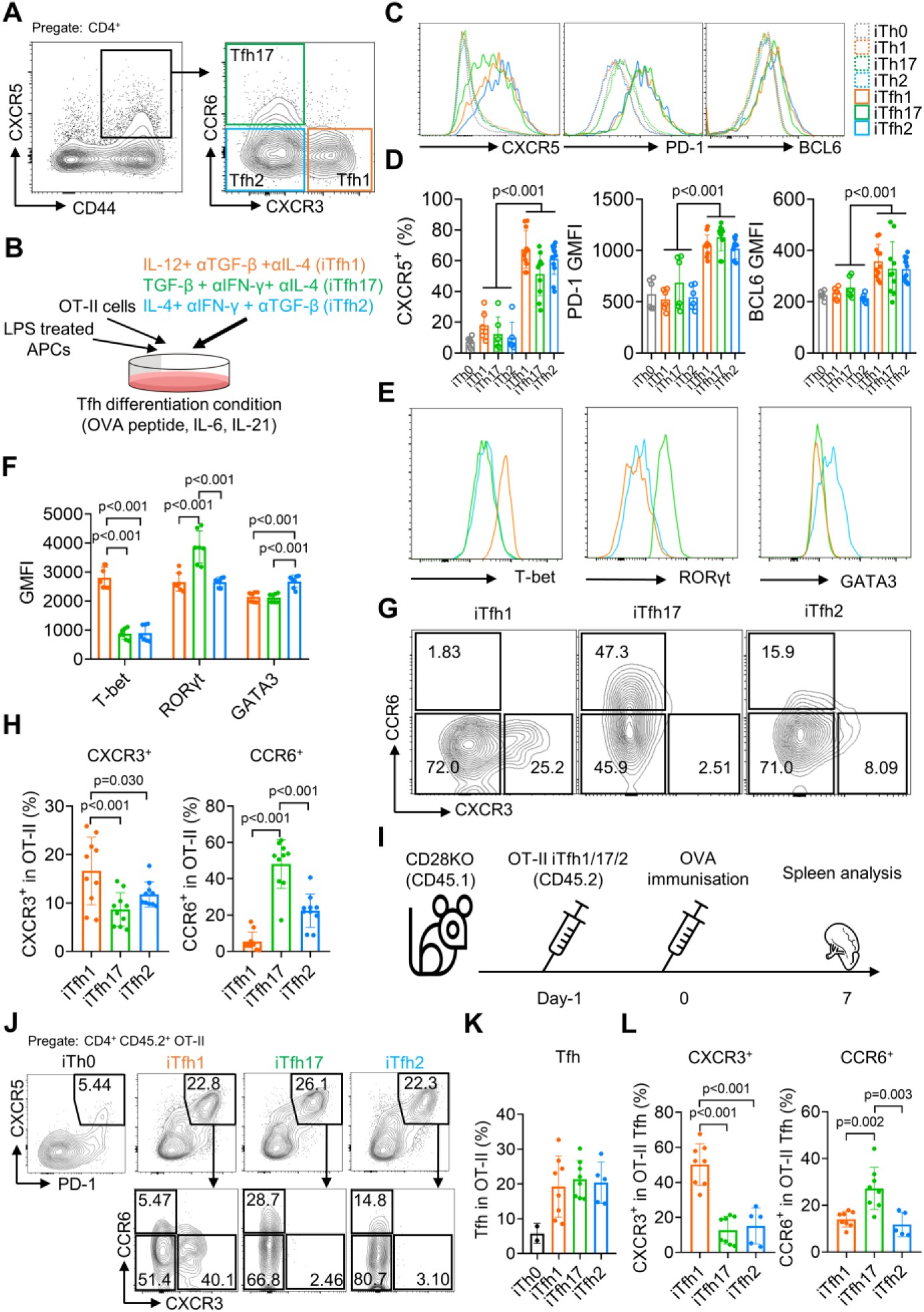
The *in vitro* differentiation of induced Tfh1, Tfh2 and Tfh17-like (iTfh1, iTfh2, iTfh17) cells. (A) Splenocytes from WT mice were analyzed and representative FACS plot for Tfh1, Tfh17 and Tfh2 cells was shown. (B-H) OT-II cells were co-cultured with WT splenocytes as antigen-presenting cells (APCs) in the presence of OVA peptide, indicated cytokines and blocking antibodies for three days before phenotypic analysis. Experiment design (B), representative FACS plots for the expression of Tfh markers CXCR5, PD-1 and BCL6 (C) transcription factors T-bet, RORγt and GATA3 (E), CXCR3 *vs* CCR6 expression (G) and statistics (D, F, H). (I-L) 5×10^4^ cultured OT-II iTh0, iTfh1, iTfh2 and iTfh17 cells were FACS-purified and separately transferred into CD28KO recipients, followed by OVA immunization in alum. The spleens were collected on day7 post-immunization for FACS analysis. Experiment design (I), representative FACS plot for Tfh percentage in OT-II cells (J), statistics of Tfh percentage in OT-II cells (K) and statistics of CXCR3/CCR6^+^ percentage in OT-II Tfh cells (L). The *p* values were calculated by two-way ANOVA for (D) and one-way ANOVA for (F, H, L). The results in (D, F, H) were pooled from 3 independent experiments. The results in (K, L) were pooled from 2 independent experiments.

To examine whether iTfh1/2/17 cells retain polarized phenotypes *in vivo*, we adoptively transferred each cell type individually into congenic CD28KO recipient mice, followed by the immunization of OVA in aluminium salt (OVA-Alum) (**Figure 1I**). After 7 days, iTfh1/2/17 cells showed the comparable ability of effector Tfh differentiation (**Figure 1J, K**) but maintain the distinction in CXCR3 and CCR6 expression aligning with their progenitors (**Figure 1J, L**). These results suggest that *in vitro* generated iTfh1/2/17 cells can be used to investigate the function of Tfh1/2/17 cells.

### iTfh17 cells are superior in memory maintenance

To compare the function of Tfh1, Tfh2 and Tfh17 *in vivo*, we adoptively transferred each of OT-II naïve T cell-derived iTfh1, iTfh2 or iTfh17 cells into congenic CD28KO recipient mice. T cells in CD28KO mice are defective in co-stimulation and unable to generate endogenous Tfh cells so antibody responses in CD28KO mice are dependent on transferred iTfh cells. After adoptive cell transfer, mice were immunized with OVA-Alum at day 0 (early immunization) or day 35 (late immunization) with the latter condition mimicking memory maintenance of Tfh1, Tfh2 and Tfh17 for extended *in vivo* resting (**Figure 2A**).

**Figure 2.**
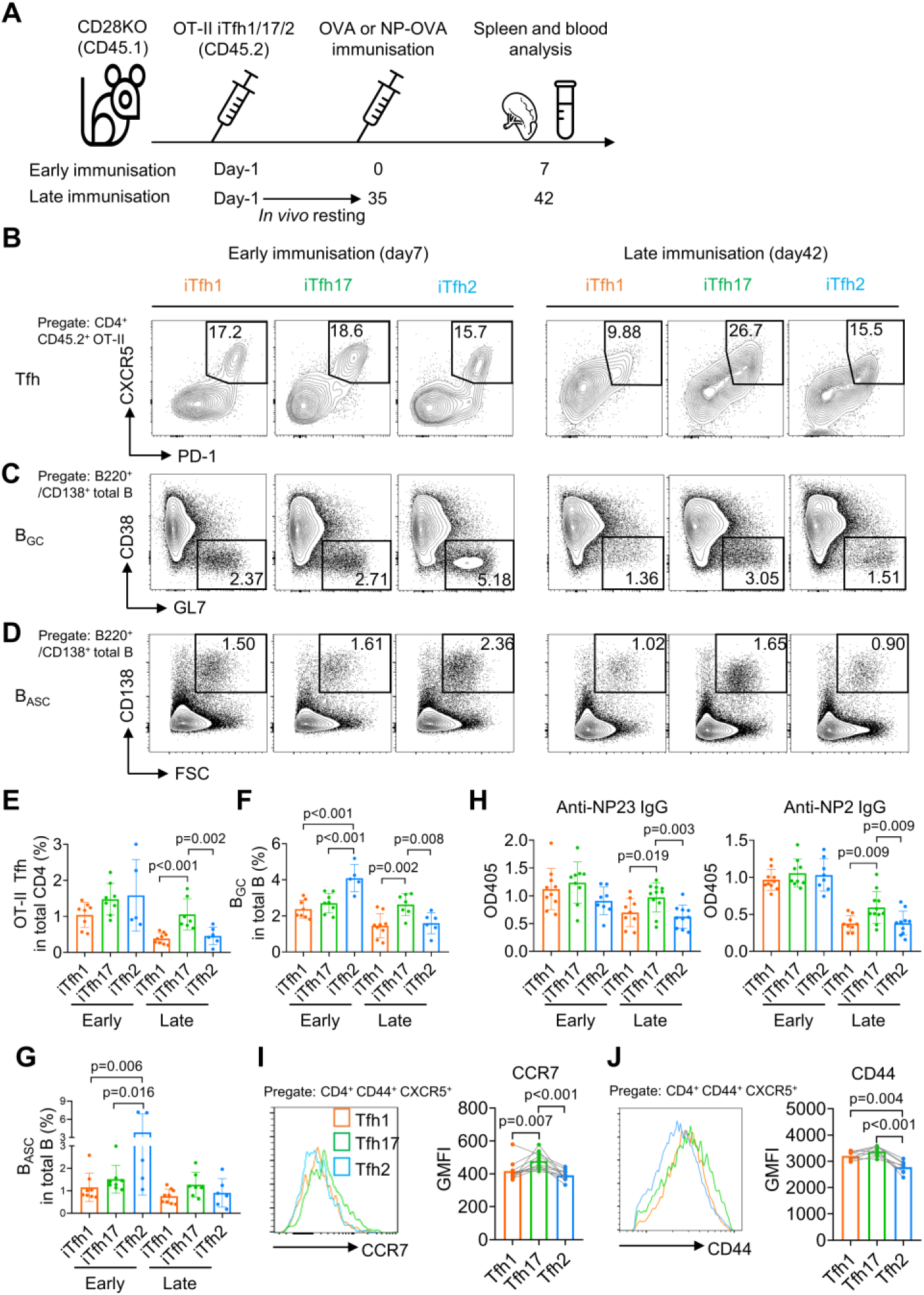
iTfh17 cells are superior in memory maintenance. (A-H) 5×10^4^ FACS-purified OT-II iTfh1, iTfh17 or iTfh2 cells were separately transferred to CD28KO recipients. The early immunization group was immunized by OVA or NP-OVA in alumn one day after the adoptive cell transfer. The late immunization group was immunized by the same antigens 35 days after the adoptive cell transfer. Spleens or serum were collected on day 7 after the immunization. Experiment design (A), representative FACS plots (B, C, D) and statistics (E, F, G) showing the percentages of Tfh cells in OT-II cells, the percentages of B_GC_ in total B cells and the percentages of B_ASC_ in total B cells. For antibody titers, statistic (H) showing OD405 values of anti- NP2 and anti-NP23 total IgG. (I-J) Tfh1/2/17 cells from mouse splenocytes were analyzed for CCR7 and CD44 expression. Representative FACS plots (I). and statistics (J) showing the expressions of CCR7 and CD44. The *p* values were calculated by one-way ANOVA. The results in (E, F, G, H, I, J) were both pooled from 2 independent experiments.

On day 7 post the early immunization, OT-II iTfh1, iTfh2 and iTfh17 cells demonstrated largely comparable Tfh differentiation and the function in supporting the generation of germinal centre B (B_GC_) cells and antibody-secreting B (B_ASC_) cells (**Figure 2B-G**). A larger magnitude of B_GC_ cells supported by iTfh2 cells might be explained by the function of IL-4 in enhancing B_GC_ generation (23). However, the outcome was very different in the scheme of late immunization whereby iTfh cells had experienced memory maintenance. After resting *in vivo* for 35 days, iTfh17 cells produced more than 2-fold of mature effector Tfh cells than those by iTfh1 or iTfh2 cells (**Figure 2B, E**), accompanied by ∼2-fold increase in B_GC_ differentiation in mice that had received iTfh17 cells than those had received iTfh1 or iTfh2 cells (**Figure 2C, F**). Although the trend of an increase in iTfh17-supported B_ASC_ differentiation didn’t reach statistical significance (**Figure 2D, 2G**), iTfh17 cells were superior to iTfh1 and iTfh2 cells in supporting the production of anti-NP IgG antibodies, after resting *in vivo* for 35 days before the immunization with antigen 4-Hydroxy-3-nitrophenyl (NP)-OVA (**Figure 2H**). Collectively, iTfh17 cells are superior to iTfh1 or iTfh2 cells in helping B cells, but only in the scheme with extended *in vivo* resting.

The selective advantage of iTfh17 cells in supporting Tfh differentiation and humoral immunity after an extended *in vivo* resting followed by immunization suggests that Tfh17 cells may outperform Tfh1 or Tfh2 cells to sustain Tfh memory. In resting mice, Tfh17 cells expressed higher CCR7 than that on Tfh1 or Tfh2 cells (**Figure 1A, 2I**), despite the highest expression of activation marker CD44 by Tfh17 cells (**Figure 2J**). CCR7^+^ marks T_CM_ cells that circulate in the blood and secondary lymphoid tissues and have longer survival and better proliferative capacity than CCR7^-^ T_EM_ cells (18, 24). We thus hypothesized that Tfh17 cells might carry certain features of T_CM_ cells suitable for memory maintenance.

### Human Tfh_CM_ and Tfh_EM_ subsets phenotypically and functionally resemble T_CM_ and T_EM_ subsets respectively

Following the observation that iTfh17 cells showed a unique advantage in memory maintenance, we set to characterize the function of human Tfh17 cells in maintaining Tfh memory. We previously reported that human cTfh cells are composed of CCR7^high^PD-1^low^ T_CM_-like and CCR7^low^PD-1^high^ T_EM_-like subsets with the latter indicating an active Tfh differentiation (4), but their function has not been formally compared. We first investigated the relationship between the two cTfh subsets and corresponding CD4^+^ T_CM_ and T_EM_ subsets by transcriptomic analysis using RNA sequencing (RNA-seq) (**Figure 3A**). As shown in an unsupervised multidimensional scaling (MDS) plot, Tfh_CM_ cells closely cluster with T_CM_ cells, and Tfh_EM_ cells fall between T_CM_ and T_EM_ cells on the major dimension1, implying that Tfh_EM_ cells are distinct from Tfh_CM_ and T_CM_ cells but also have effector programs different from T_EM_ cells (**Figure 3B**). Previous studies reported that cTfh cells predominantly show CCR7^+^ CM phenotype (25). Our transcriptomic analysis indeed suggests that Tfh_CM_ cells acquire a quiescent state hardly distinguishable from T_CM_ cells. In contrast, Tfh_EM_ cells’ transcriptomes are clearly separated from those of T_EM_ cells, presumably caused by the divergent effector function of Tfh cells as compared to other effector Th1, Th2 or Th17 cells. In line with this, the top 50 differentially expressed genes (DEG) indicate effector genes such as *ZEB2* and *TBX21* (26) were highly expressed in T_EM_ cells, intermediate levels in Tfh_EM_ cells, and lowest in Tfh_CM_ and T_CM_ cells (**Figure 3C**). In top 50 hallmark gene sets identified by gene set enrichment analysis (GSEA) between Tfh_EM_ *vs* Tfh_CM_ cells or T_EM_ *vs* T_CM_ cells, 37 gene sets were significantly enriched by both comparisons (NES discrepancy > 2, **Figure 3D**), suggesting that the transcriptomic features and regulation between Tfh_EM_ *vs* Tfh_CM_ cells are overall similar to those between T_EM_ *vs* T_CM_. Despite T_EM_ and Tfh_Em_ cells show distinct transcriptomes and locate separately in the MDS plot (**Figure 3B**), the key gene sets that are related to common effector T cell function (activation, effector differentiation and cell cycle entry) were both positively enriched in comparisons between Tfh_EM_ *vs* Tfh_CM_ cells or T_EM_ *vs* T_CM_ cells (**Figure 3E**). Therefore, Tfh_CM_ and Tfh_EM_ cells resemble T_CM_ and T_EM_ cells respectively at the transcriptomic levels.

**Figure 3.**
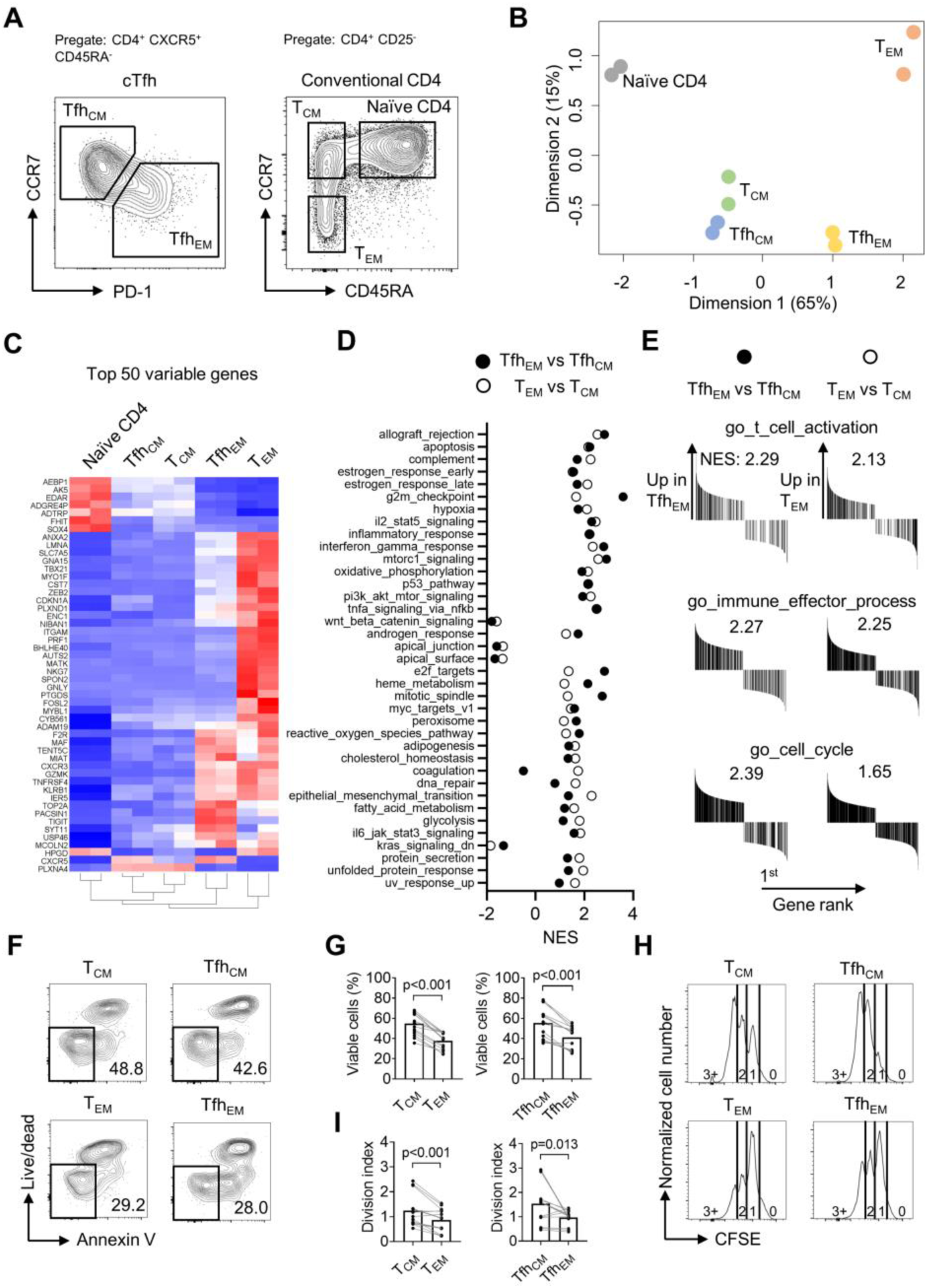
Human Tfh_CM_ and Tfh_EM_ subsets phenotypically and functionally resemble T_CM_ and T_EM_ subsets respectively. (A-E) Naïve, T_CM_, T_EM_, Tfh_CM_ and Tfh_EM_ cells were FACS-purified from PBMC of two healthy donors and bulk RNA-seq was performed for differentially expressed genes analysis and gene set enrichment analysis (GSEA). (A) Representative FACS plot showing the gating strategy for indicated subsets. (B) MDS plot showing sample distribution. (C) Heatmap of the top 50 variable genes normalized by z-score. (D) Summarized normalized enrichment score (NES) of significantly enriched (*p* <0.05, FDR < 0.25) hallmark gene sets by either Tfh_EM_ *vs* Tfh_CM_ or T_EM_ *vs* T_CM_. (E) GSEA on selected gene sets were performed on Tfh_EM_ *vs* Tfh_CM_ and T_EM_ *vs* T_CM_ and the number indicates NES. (F-G) FACS-purified T_CM_, T_EM_, Tfh_CM_ and Tfh_EM_ cells were rested in complete RPMI for 3 days. Representative FACS plots (F) and statistics (G) showing the percentages of viable cells. (H-I) FACS-purified T_CM_, T_EM_, Tfh_CM_ and Tfh_EM_ cells were labelled with CFSE and stimulated by anti-CD3/CD28 for 2.5 days. Representative FACS plots (H) and statistics (I) showing the CFSE fluorescence intensity and the division index. The *p* values were calculated by Wilcoxon matched-pairs signed-rank test. The results in (G, I) were pooled from 5 healthy individuals with each conducted in 3 technical replicates.

We next compared Tfh_CM_ and Tfh_EM_ subsets for survival and stimulation-induced proliferation in culture, which were applied to characterize the difference between T_CM_ and T_EM_ cells (18). In non-stimulation culture for 3 days, T_CM_ and Tfh_CM_ cells retained ∼50% viability while T_EM_ and Tfh_EM_ cells showed poorer survival of ∼30% (**Figure 3F, G**). To measure the proliferative potential, all subsets were labeled with carboxyfluorescein succinimidyl ester (CFSE) and stimulated by anti-CD3/CD28 for 2.5 days. While the majority of T_EM_ or Tfh_EM_ cells underwent division once, most T_CM_ or Tfh_CM_ cells reached the second or third division, indicating a better proliferative potential (**Figure 3H, I**). Collectively, CCR7^high^PD-1^low^ Tfh_CM_ and CCR7^low^PD-1^high^ Tfh_EM_ subsets showed not only transcriptomic profiles resembling their counterpart T_CM_ and T_EM_ cells but also functional characteristics of survival and proliferative capacity (18, 24).

### Human Tfh_CM_ cells are enriched with the Tfh17 subset whereas Tfh_EM_ cells are enriched with the Tfh1 subset

From a cohort of healthy donors (*N* = 33, **Table S1**), we analyzed CCR7^high^PD-1^low^ Tfh_CM_ and CCR7^low^PD-1^high^ Tfh_EM_ cells for the percentages of Tfh1/2/17 subsets based on CXCR3 and CCR6 expression. In agreement with the higher expression of CCR7 on mouse Tfh17 than that on Tfh1 or Tfh2 cells (**Figure 2I**), human Tfh_CM_ cells were dominated by the Tfh17 subset (mean = 51.44%), followed by the Tfh2 subset (mean = 16.19%) and the Tfh1 subset (mean = 12.17%) (**Figure 4A**), whereas Tfh_EM_ cells were dominated by the Tfh1 subset (mean = 34.06%,) (**Figure 4B**). The population of Tfh cells that expresses both CXCR3 and CCR6 has been reported and also presented in our samples. Due to the fact that CXCR3^+^CCR6^+^ cTfh cells were fewer than Tfh1, Tfh2 or Tfh17 cells and their ontogeny remains to be fully revealed (10), we did not include this population in the following analyses. To avoid the influence of individual variation of Tfh1/2/17 polarization due to different histories of immune exposure and examine whether there is an intrinsic difference of Tfh1/2/17 frequencies between Tfh_CM_ or Tfh_EM_ cells, the percentages of Tfh1/2/17 cells in Tfh_CM_ cells were normalized to those in Tfh_EM_ cells in each individual, which demonstrated the highest Tfh_CM_/Tfh_EM_ ratio for Tfh17 (mean = 4.18), an intermediate ratio for Tfh2 (mean = 3.38), and the lowest ratio for Tfh1 (mean = 0.75) (**Figure 4C**). The highest ratio (>> 1) for Tfh17 indicates Tfh_CM_ cells are highly enriched with the Tfh17 subset whereas the lowest ratio (< 1) for Tfh1 indicates that Tfh_EM_ cells are enriched with the Tfh1 subset. We also measured the expression of hallmark transcription factors *TBX21*, *GATA3* and *RORC* and cytokines IFN-γ, IL-4 and IL-17A that are selectively expressed in Tfh1/2/17 cells respectively (10). These molecules for effector Th functions are abundantly expressed by T_EM_ cells but are downregulated in T_CM_ cells which enter into a resting state (18). Indeed, the expression of effector transcription factors and cytokines was consistently lower in Tfh_CM_ cells than those in Tfh_EM_ cells (**Figure 4D, F**). Notably, the ratios of expression (Tfh_CM_/Tfh_EM_) demonstrate modest reductions of 20-40% in Tfh17 related markers RORγt and IL-17A, in contrast to vast reductions of 70-80% in Tfh1 related markers T-bet and IFN-γ (**Figure 4E, G**). Such results of transcription factor and cytokine expression support the conclusion for an enrichment of the Tfh17 subset and a loss of the Tfh1 subset in Tfh_CM_ cells.

**Figure 4.**
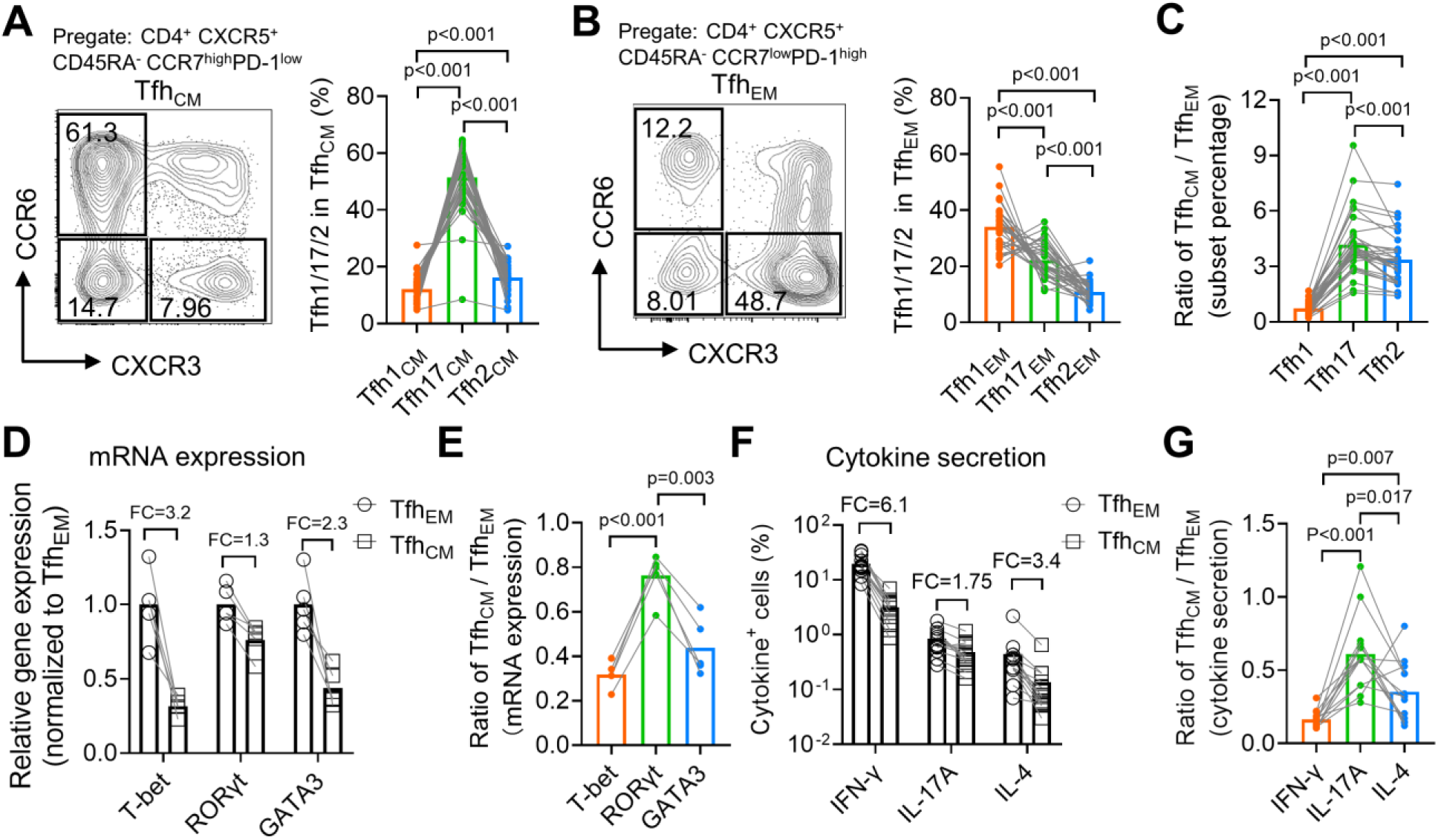
Human Tfh_CM_ cells are enriched with the Tfh17 subset whereas Tfh_EM_ cells are enriched with the Tfh1 subset. (A-C) Human PBMC samples from 33 healthy blood donors were analyzed. Representative FACS plots and statistics showing the percentages of Tfh1, Tfh2 and Tfh17 cells in Tfh_CM_ (A) or Tfh_EM_ (B) subsets. Tfh_CM/_Tfh_EM_ ratios for Tfh1/2/17 in each individual were calculated (C). (D-E) FACS-purified Tfh_EM_ and Tfh_CM_ from 5 healthy individuals were analyzed for the expressions of indicated transcription factors by qPCR. The statistics for relative gene expression 2^-ΔΔCt^ (normalized to Tfh_EM_) (D) and Tfh_CM_/Tfh_EM_ ratios (E). (F-G) PBMC from 13 healthy individuals were analyzed for the secretions for indicated cytokines post PMA/ionomycin stimulation. The statistics for the percentages of cytokine^+^ cells (F) and the Tfh_CM_/Tfh_EM_ ratios (G). FC: average fold change. The *p* values were calculated by Friedman test.

### HBV antigen-specific Tfh17 cells are preferentially maintained in memory phase

Tfh_EM_ to Tfh_CM_ phenotype conversion occurs over the period of a few weeks when the antigen is absent (4). The enrichment of the Tfh17 subset in human Tfh_CM_ cells suggest the Tfh17 subset in Tfh_EM_ cells may persist longer than the Tfh1 or Tfh2 subsets. The phenomenon could also result from a biased Tfh_CM_ phenotype of Tfh17 cells generated even early in immune responses. To tease apart the cause, we next examined the phenotype of antigen-specific human cTfh cells over a period that expands both effector and memory phases after vaccination or infection.

Childhood HBV vaccination doesn’t always provide life-long protection with a proportion of vaccinees with antibody titers at an undetectable level in adulthood (27). HBV boosting vaccination is recommended for high-risk populations such as medical practitioners including medical students in China. A cohort of medical students (*N* = 38) with serum negative for anti-HBV surface antigen (HBVSA) antibody were recruited (**Table S1**). Peripheral blood mononuclear cells (PBMCs) were collected at day 7 before and day 7 and 28 after the immunization (**Figure 5A**). Antigen-induced marker (AIM) assay was used to examine HBV vaccine-specific T cells by culturing PBMCs with HBVSA for 18h and detecting the PD- L1^+^OX40^+^CD25^+^ cells as the antigen-specific population (**Figure 5B**). This method has been applied to characterize antigen-specific cTfh cells (28, 29). Such stimulation did not change the expression of CXCR3 and CCR6 on cTfh cells **(Figure S2A-C)**, indicating that AIM assays are suitable to characterize antigen-specific Tfh1/2/17 cells. As reported (29), a background in AIM assays exists in a small proportion of samples whereby PD-L1^+^OX40^+^CD25^+^ cells were detected in control cultures without antigen stimulation (**Figure S2D**). To specifically quantify antigen-specific response, we subtracted the value of antigen-stimulation culture by the background value from the control culture without antigen stimulation. Normalized values were then used to calculate the percentages of Tfh1, Tfh2 and Tfh17 cells (**Figure S2E**).

**Figure 5.**
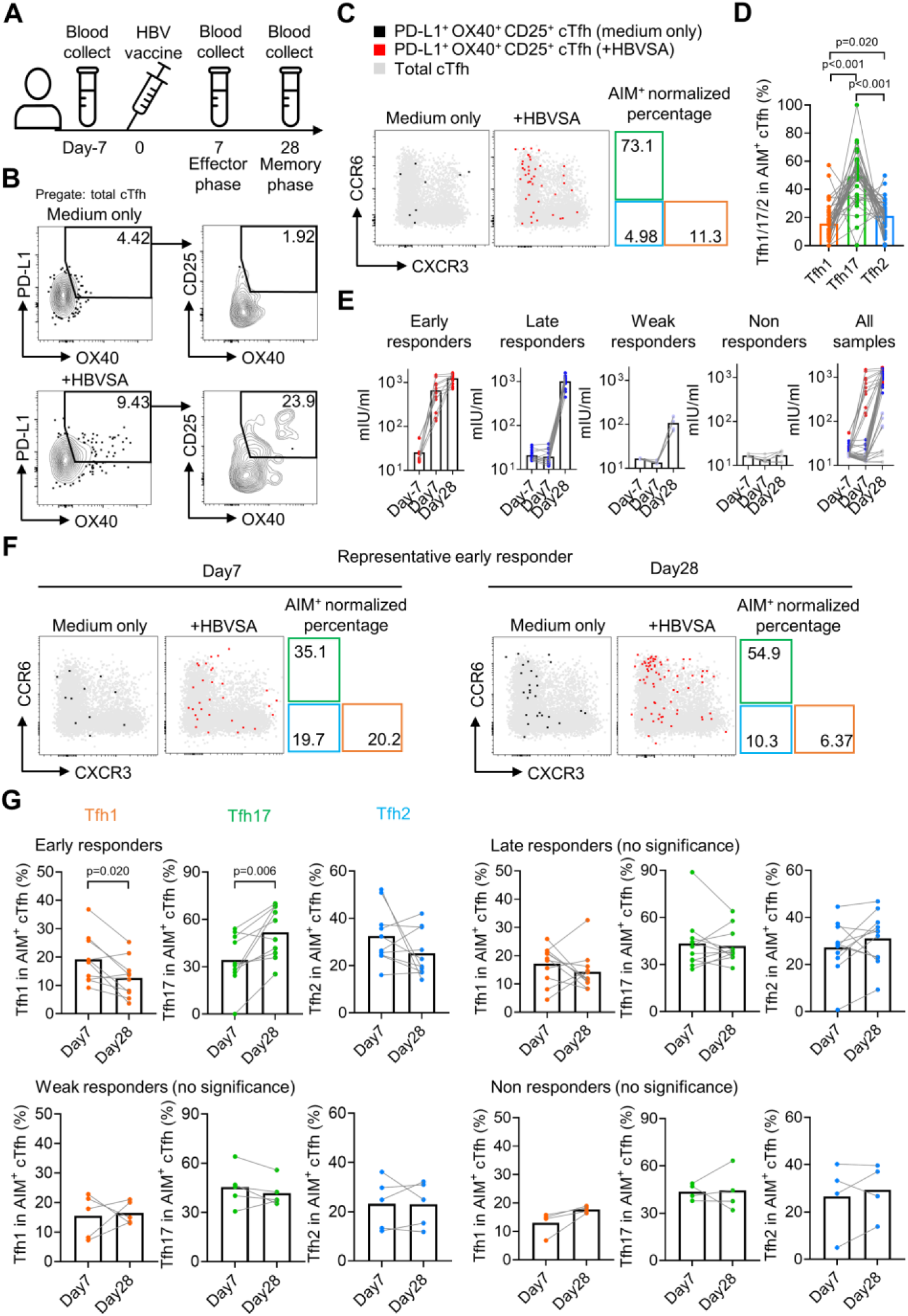
HBV antigen-specific Tfh17 cells are preferentially maintained in memory phase. Blood samples from HBV vaccinated healthy individuals (*N* = 38) were collected on indicated time points before/after HBV vaccination, and serum was diluted 10 times to analyse the anti-HBVSA antibody titer by ELISA. PBMC were also isolated and cultured with or without 20 µg/mL HBVSA for 18h, followed by FACS to analyse the phenotype of HBVSA-specific cTfh cells. Experiment design (A) and representative FACS plot (B) showing the gating strategy to detect HBVSA-specific cTfh cells by AIM assay. Representative FACS plot (C) and statistics (D) showing the percentage of Tfh1/2/17 cells in HBVSA-specific cTfh cells before vaccination. Classification (E) of 38 individuals into 4 groups was based on their anti-HBVSA antibody titers. Representative FACS plot (F) for an early responder showing the percentage of Tfh1/2/17 cells in HBVSA-specific cTfh. Statistics (G) showing the percentage of Tfh1/2/17 cells in HBVSA-specific cTfh on day 7 and day 28 after the vaccination in all defined groups (*N* = 30, 8 samples with poor signals in AIM assay were excluded). The *p* values were calculated by Wilcoxon matched-pairs signed-rank test.

In all subjects negative for HBVSA antigen and anti-HBVSA antibody, the average percentage of the Tfh17 subset in HBVSA-specific memory cTfh cells was 49.31%, whereas the average percentage of the Tfh1 subset was 15.7% (**Figure 5C, D**). In alignment with the results on total Tfh_CM_ cells (**Figure 4**), HBVSA-specific Tfh_CM_ cells were also enriched with the Tfh17 subset.

To tease apart whether the Tfh17 enrichment was caused by the biased generation or better maintenance, we then analysed PBMC samples collected at day 7 and day 28 after vaccination (**Figure 5A**). The cohort was divided into 4 groups based on vaccine responses measured by antibody titers (**Figure S3A**): early responders (titer > 100 mIU at day7, titer = 1248±324.4 mIU at day 28, *N* = 11), late responders (titer < 100 mIU at day 7 and > 200 mIU at day 28, titer = 997.1±289.7 mIU/ml at day 28, *N* = 15), weak responders (50 mIU < titer < 200 mIU at day 28, titer = 107.9±37.2 mIU at day 28, *N* = 6) and non-responders (titer < 50 mIU on day 28, titer = 17.28±4.379 mIU at day 28, *N* =6) (**Figure 5E**). Tfh activation measured by a trend of increase in HBVSA-specific Tfh_EM_ cells by (2.84-fold, day -7 v.s. day 7, *P*-value = 0.074, not reaching statistical significance) was observed only in early responders but no other groups (**Figure S3B, C).** We next focused on the kinetics of HBVSA-specific cTfh subsets in early responders. At day 7 post vaccination, the percentages of three subsets in HBVSA-specific cTfh cells ranged from 20% to 30% and showed no significant difference (**Figure S3D**). Of note, the percentages of the Tfh17 subset significantly increased from ∼30% to ∼50% from day 7 to 28 (*P*-value = 0.006). In contrast, the percentages of the Tfh1 subset dropped significantly from ∼20% to ∼10% (*P*-value = 0.020). The percentages of Tfh2 remained largely unchanged (**Figure 5F, G**). As a result, Tfh17 cells dominated HBVSA-specific cTfh cells in the memory phase of day 28 post vaccination **(Figure S3D)**. Therefore, the Tfh17 enrichment in Tfh_CM_ cells results from an advantage of Tfh17 cells in memory maintenance, rather than a biased phenotype to Tfh_CM_ cells. Significant changes in Tfh1 and Tfh17 percentages from day 7 to day 28 were selective in early responder group but not in three other groups (**Figure 5G**), and only observed in HBVSA- specific cTfh cells but not in total cTfh in early responders (**Figure S3E**), suggesting that the dynamic changes were specific to HBVSA-specific cTfh response.

### Influenza virus-specific cTfh cells show Tfh1 signatures in effector phase but Tfh17 signatures in memory phase

Single-cell RNA-seq (scRNA-seq) paired with TCR sequencing facilitates the characterization of the phenotype and function of antigen-specific T cell clones in an immune response. We took the advantage of this new technology to analyze the characteristics of influenza haemagglutinin (HA)-specific CD4^+^ T clones in a published dataset of scRNA/TCR-seq from four healthy individuals with influenza vaccination (30). We compared HA-specific T cell clones before the vaccination (memory phase) and day 12 after the vaccination (effector phase) (**Figure 6A**). HA-specific CD4^+^ T cells before and after the vaccination were pooled to generate unsupervised clustering, in which CXCR5-expressing clusters 2-5 were enriched of cTfh cells, in which a total of twelve major CD4^+^ T clones (clonal abundance ≥ 10) were identified (**Figure 6B**). To investigate Tfh subsets-associated features in HA-specific clonal cTfh cells, we applied Tfh1 or Tfh17 signature gene sets derived from bulk RNA-seq for Tfh1/2/17 cells **(Figure S4A)** (31) in clonal cTfh cells. The scores of Tfh1 signature were higher in clonal cTfh cells in the effector phase than those in the memory phase (*P*-value = 0.041); by contrast, the scores of Tfh17 signature were lower in the effector phase than in the memory phase (*P*-value = 0.002) (**Figure 6C, D**). The divergence of Tfh1 and Tfh17 signatures between the effector and memory phases was consistent in individual donors **(Figure S4B)** and at the level measured by twelve major clones (**Figure 6E, F**). Therefore, we conclude that the advantage of Tfh17 cells in memory maintenance is consistently observed among different cohorts with different types of vaccines.

**Figure 6.**
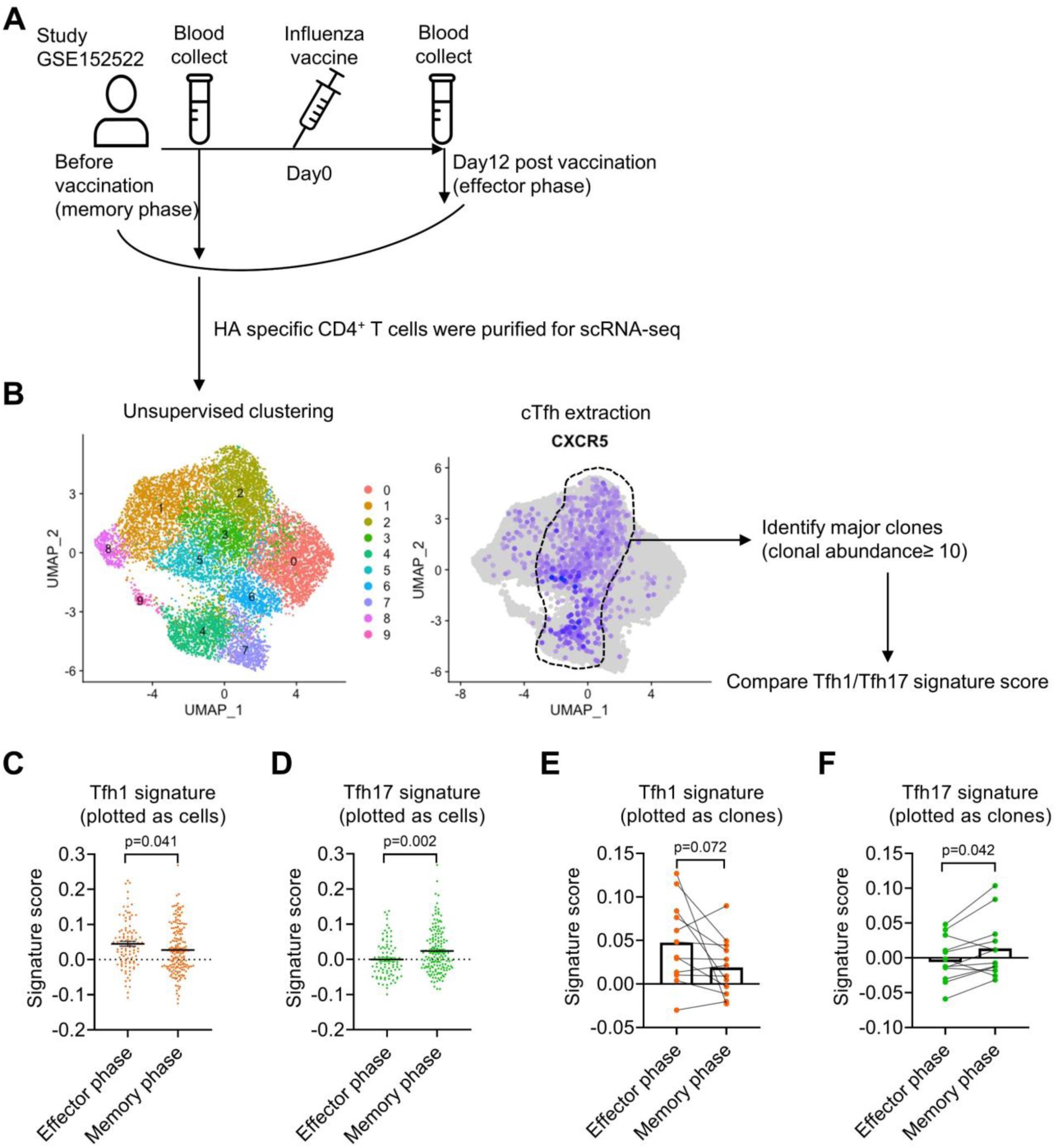
Influenza virus-specific cTfh cells show Tfh1 signatures in effector phase but Tfh17 signatures in memory phase. The single-cell RNA-seq dataset (GSE152522, the experiment design (A)) was analyzed to identify CXCR5- expressing cTfh clusters (B), which contain 12 major clones with a total of 249 cells. Comparison of Tfh1 and Tfh17 signature scores between effector and memory phase cTfh cells based on each cell or clone were shown in (C, D) and (E, F). The signature score of each clone was calculated as the mean value of the signature scores of all the cells in this clone. The *p* values were calculated by unpaired *t*-tests for (C, D) and paired *t-*tests for (E, F).

### Tfh17 cells are long-lived and accumulate with aging

Our previous experiments have demonstrated that antigen-specific mouse iTfh17 cells and vaccine-specific human Tfh17 cells are superior to Tfh1 and Tfh2 cells in maintaining Tfh memory for a period of about one month (HBV) or less than one year (influenza vaccine). We next ask whether Tfh17 cells can persist for even longer periods, such as years. In a cohort of adults (*N* =20, **Table S1**), we examined cTfh cells specific to vaccines for tetanus toxoid and measles, both administrated in childhood (**Figure 7A**). Given that community transmission of tetanus and measles is very rare (32), cTfh cells specific to tetanus toxoid and measles in adults were likely induced many years ago by childhood vaccination (33–35). The average percentages of the Tfh17 subset in vaccine-specific memory cTfh cells were 55.22% and 45.07% for tetanus toxoid and measles respectively, which were more than twofold higher than the Tfh2 percentages and more than threefold higher than the Tfh1 percentages (**Figure 7B, C**). These results suggest Tfh17 cells may maintain Tfh memory for more than a decade. We also asked whether the Tfh17 dominance in memory phase was also applied to cTfh cells induced by SARS-CoV-2 infection. We examined convalescent patients with Covid-19 showing SARS-CoV-2-specific IgG antibodies (*N* = 13, **Figure S5A** and **Table S1**). Similar to vaccine-specific cTfh cells, the Tfh17 percentages in SARS-CoV-2-specific cTfh cells were much higher than the Tfh1 or Tfh2 percentages (mean, 59.03% v.s. 12.87% or 7.73%) (**Figure S5B, C**).

**Figure 7.**
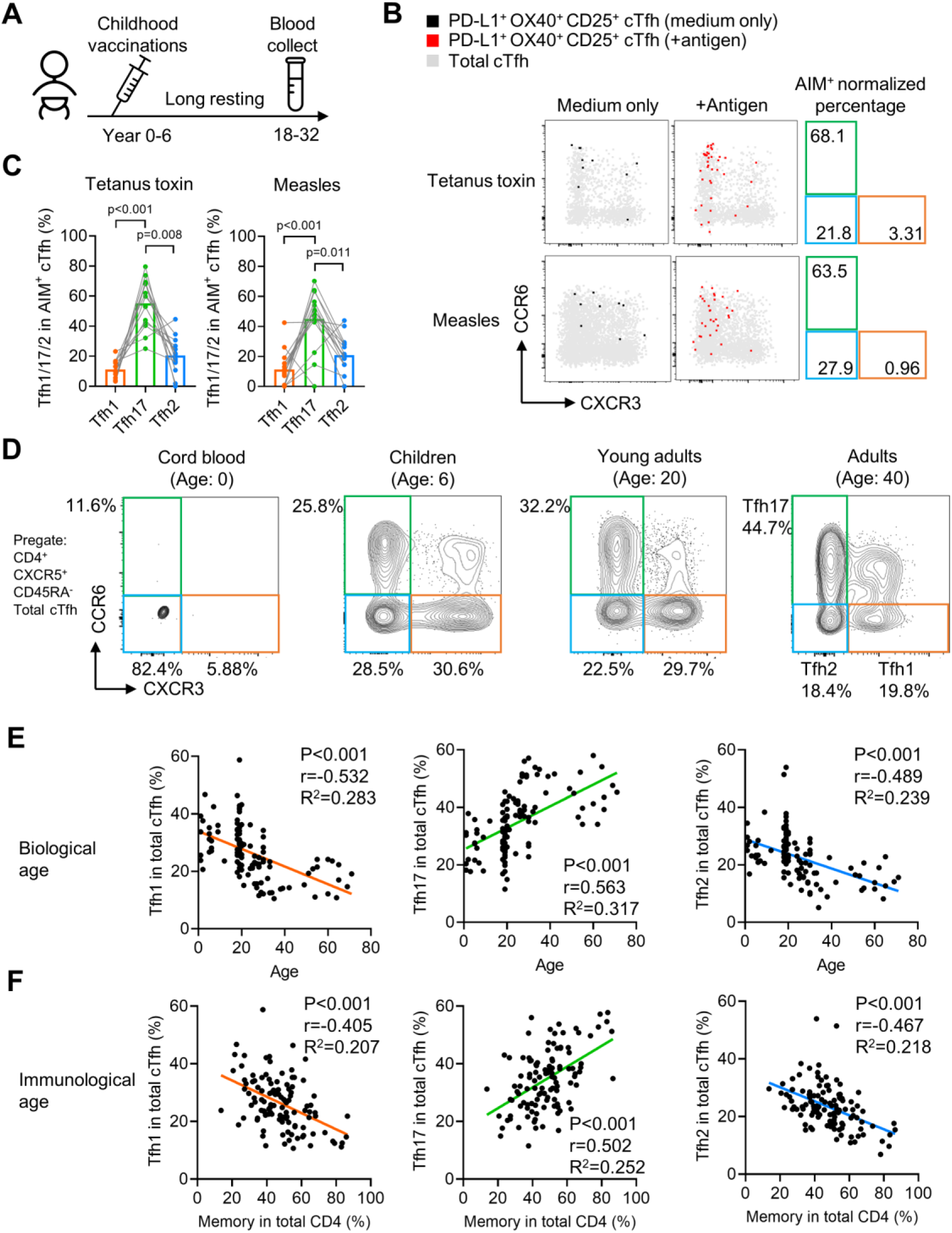
Tfh17 cells are long-lived and accumulate with aging. (A-C) PBMC samples from 20 healthy individuals were cultured for 18h with or without indicated antigens, followed by FACS to detect the phenotype of antigen-specific cTfh cells. Experiment design (A), representative FACS plot (B) and statistics (C) showing the percentage of Tfh1/2/17 cells in antigen-specific cTfh cells against tetanus toxin or measles. (D-F) PBMC samples from individuals of different ages were analysed. Representative FACS plots (D) showing the percentages of Tfh1/2/17 cells in total cTfh cells in individuals of different ages. Correlations tests between the biological age (E) or immunological age (F) with the percentages of Tfh1/2/17 cells in total cTfh cells. Cord blood samples were excluded from the correlation tests because of insufficient cTfh cell numbers. The *p* values were calculated by Friedman test for (C) and Pearson correlation for (E) and (F).

If antigen-specific Tfh17 cells are long-lived and superior to Tfh1 and Tfh2 cells for persistence, we would expect a preferential accumulation of Tfh17 cells over Tfh1 or Tfh2 cells along with aging. By pooling results of cTfh characterization from cord blood, children, young and middle-aged adults and the elderly (**Table S1**), we indeed observed that the Tfh17 percentages in total cTfh cells positively correlated with biological ages whereas the percentages of Tfh1 or Tfh2 subsets showed negative correlations (*P*-value < 0.001) (**Figure 7D, E**). In addition to ‘biological ages’, individuals also differ in ‘immunological ages’ due to various antigen exposure histories.

We used the percentages of CD45RA^-^ memory cells in total CD4^+^ T cells as the surrogates for ‘immunological ages’, and again found a strong positive correlation between immunological ages and the Tfh17 percentages in total cTfh cells, in contrast to the negative correlations with the Tfh1 or Tfh2 percentages (*P*-value < 0.001) (**Figure 7F**).

In conclusion, Tfh17 cells are superior to Tfh1 and Tfh2 cells in Tfh memory maintenance, a phenomenon consistently observed in vaccination, infection and natural antigen exposure.

## Discussion

Morita et al. reported circulating Tfh memory cells comprise different subsets related to Th1, Th2 and Th17 cells (10), which has advanced our understanding of Tfh subsets and helped to delineate the relationship between Tfh and other Th subsets. While the signatures of Tfh2 and Tfh17 activation were commonly reported in allergic and autoimmune diseases (36, 37), the activation of Tfh1 cells was a prominent feature and associated with pathogen-specific antibody production in influenza vaccination (15) and infections by HIV (38), malaria (14) and more recently SARS-CoV-2 (39, 40). Beyond their distinct function in mediating isotype class switching (41, 42), other difference between these Tfh subsets remains largely unknown, possibly due to the experimental hurdle of a low frequency of Tfh subsets in human blood and the lack of culture method for *in vitro* Tfh subset generation and *in vivo* functional characterization.

We adopted the published method for the *in vitro* induction of antigen-specific Tfh differentiation (19) and modified the protocol by adding the conditions biased for Th1/2/17 polarization. This method corroborated the report that Tfh cells are plastic and carry positive epigenetic markings for Th1/2/17 cells (21). Notably, for the iTfh17 condition (0.1 ng/mL TGF- β + 100 ng/mL IL-6 + 50 ng/mL IL-21), TGF-β were used in a concentration of 0.1 ng/mL, much lower than those for Th17 and Treg polarization (normally 1-10 ng/mL). The condition successfully generated iTfh17 expressing both CXCR5, PD-1, BCL6 and RORγt. This phenomenon might also help to reconcile the reports that TGF-β signaling can either inhibit or support Tfh differentiation (43, 44), probably determined by the TGF-β signal strength.

The generation of antigen-specific iTfh1/2/17 cells in decent numbers using this modified method allowed us, for the first time, to compare Tfh1, Tfh2 and Tfh17 function *in vivo*. The results from the adoptive transfer experiment revealed that, after an extended period of *in vivo* resting to mimic memory maintenance, iTfh17 cells showed a better function than iTfh1/2 cells in supporting humoral immunity. In agreement with this, the Tfh17 subset in human cTfh cells also showed superiority over Tfh1 and Tfh2 subsets for memory maintenance. Tfh17 cells predominantly showed the Tfh_CM_ phenotype as long-lived memory cells and dominated the long-lived pool of antigen-specific memory Tfh cells for vaccines of HBV, influenza, tetanus and measles. In contrast, the human Tfh1 subset appears short-lived and accounted for the least proportion of long-lived antigen-specific memory Tfh cells. Tfh1 is the major Tfh subset induced by influenza infection and vaccination (15). The short-lived characteristics of Tfh1 cells may partially contribute to a relatively short period of humoral immunity after influenza vaccination (45). In SARS-CoV-2 infection, patients with acute infection demonstrated a Tfh1-biased profile, while convalescent patients increased the proportions of virus-specific Tfh17 cells, again supporting the notion that Tfh17 cells represent the population better in memory maintenance (39, 46). In malaria infection, a recent study also reported Tfh17 and Tfh2 cells, rather than Tfh1 cells showed a Tfh_CM_ phenotype (47). All such evidence suggests the superiority of Tfh17 subset in memory maintenance appears to be a common feature for immune responses induced by both vaccination and infection. It should be noted that our results should not be misinterpreted as that Tfh17 cells are always the major subset for Tfh memory cells. In the case of SARS-CoV-2 mRNA vaccine which induces strong Th1-polarized response, Tfh17 cells are essentially not induced and the Tfh memory are maintained in the absence of Tfh17 cells (48–50).

The mechanism underlying the advantage of Tfh17 cells in memory maintenance is an area we will focus on in the following studies. Intriguingly, Th17 cells were reported to have “stem cell-like” features and are long-lived (51–53). The proposed mechanisms are diverse, which include a high expression of Tcf1 in Th17 cells, a key transcription factor that regulates T cell memory generation and self-renewal and favorable expression of anti-apoptosis Bcl-2 family genes to sustain the longevity (52, 53). The Th17’s hallmark transcription factor RORγ has been shown to directly promote T cell survival by enhancing Bcl-xL expression (54). Effector and memory Tfh cells are critically regulated by specific cell death pathways of ferroptosis and pyroptosis (55, 56). Future works are required to test whether these mechanisms also apply in the survival and self-renewal of Tfh17 cells.

Unveiling the superiority of Tfh17 in Tfh memory maintenance can help us to improve the rationale-based vaccine development. Many vaccines, including conventional vaccines for influenza virus and novel mRNA vaccines for SARS-CoV-2, induce Th1 responses and Tfh1- associated humoral immunity (15, 57, 58). According to our results, Tfh1 cells are short-lived, which might curb the duration of vaccine-mediated protection. New strategies might be taken to direct vaccination for more Tfh17 induction which can support a better Tfh memory formation and potentially prolong vaccine protection.

## Materials and Methods

### Study design

This study aims to investigate the memory function of different Tfh subsets in human and mice. Tfh1/2/17 subsets in total and antigen-specific cTfh cells from healthy donors and vaccinees were analysed for phenotypes and kinetics. *In vitro* generated Tfh1/2/17 (iTfh1/2/17) cells were analysed for phenotypes and also function after being transferred into recipient mice followed by immunizations. Human cohort samples sizes varied and were guided by previous studies. Mouse sample sizes of three to five per group per time point were used for experiments to detect significant differences between groups while minimizing the use of laboratory animals. Mice were randomly assigned, age and gender matched between groups. The investigators were blinded in collecting raw data from human and mouse samples.

### Human samples

Demographics of human samples were shown in Table S1. Written informed consent was obtained from participants or the parents of children participants according to the ethics approved by human ethics committees of Renji Hospital affiliated to Shanghai Jiao Tong University School of Medicine (KY2019-161), Fourth Military Medical University (KY20163344-1), Tongji Hospital (NCT05009134), Shanghai Children’s Medical Centre affiliated to Shanghai Jiao Tong University School of Medicine and Obstetrics and Gynecology Hospital of Fudan University (Kyy2018-6). Whole blood samples from healthy individuals (cTfh phenotyping, *N* = 33; Measles and TT AIM assay, *N* = 20) were collected from Renji Hospital affiliated to Shanghai Jiao Tong University School of Medicine, Shanghai, China. Whole blood samples from healthy volunteers (*N* = 38) who received the standard recombinant HBV vaccine (Shenzhen Kangtai Biological Products Co.) were recruited by Fourth Military Medical University, Xi’an, China. Whole blood samples from healthy volunteers (qPCR and cytokine assay, *N =* 14; Recovered Covid-19 patients, *N =* 13) were collected from Tongji Hospital affiliated to Huazhong University of Science and Technology Tongji Medical College, Wuhan, China. Whole blood samples from children (*N* = 18) were collected from Shanghai Children’s Medical Centre affiliated to Shanghai Jiao Tong University School of Medicine, Shanghai, China. Cord blood samples (*N* = 5) were collected from Obstetrics and Gynecology Hospital of Fudan University, Shanghai, China. Buffy coats from healthy donors for bulk RNA-seq were obtained from the blood bank of Changhai Hospital affiliated to Navy Medical University, Shanghai, China.

### Mice

CD45.1 WT, CD45.2 WT, CD28KO and OT-II mice were maintained on a C57BL/6 background and housed in specific pathogen-free conditions in the Australian Phenomics Facility (APF). All animal experiments were carried under protocols (ethics number: A2019/36) approved by ANU’s animal ethics committee.

### PBMC and plasma isolation

Blood from human and mouse were collected in BD Vacutainer Blood Collection Tubes. After centrifugation (400 g, 20°C, 5 min), plasma was collected and stored in -80°C for further analysis. Blood cells or buffy coats were diluted in PBS and gently loaded onto the Ficoll-Paque Plus (GE Healthcare) at the volume ratio of 1:1, followed by density gradient centrifugation (450 g, 20°C, 20 min, no brake). PBMC were then aspirated and resuspended in cold PBS for further experiment.

### Antigen induced marker assay (AIM assay)

Cryopreserved PBMC were thawed, washed and counted. 5 × 10^5^ PBMCs were resuspended in 200 µL complete RPMI media (3% FBS for AIM assay) and cultured in a 96-well flat-bottom plate for 18h in the presence of 20 µg/mL recombinant HBVSA (Beijing Bioforce), tetanus toxin (Sigma), measles (GenWay) or 1 µg/mL SARS-CoV-2 Prot_S (Miltenyi Biotec), no antigen was added to control wells. At least 6 wells were seeded (3 antigen treated wells + 3 medium only wells) for each PBMC sample. FACS was performed and the replicates for each sample were merged for downstream analysis.

### Quantitative RT-PCR

Total RNA was extracted from sorted T cell subsets using Trizol reagent (Thermo Fisher Scientific) and reverse-transcribed to cDNA using PrimeScript RT reagent kit (TaKaRa Biotechnology). RT-PCR was performed with StepOnePlus (Applied Biosystems) using SYBR Green PCR Master Mix (Thermo Fisher Scientific) with specific primers. The following primers (5’->3’) were used. *TBX21*: forward, CACTACAGGATGTTTGTGGACGTG, reverse, CCCCTTGTTGTTTGTGAGCTTTAG; *GATA3*: forward, TGTCTGCAGCCAGGAGAGC, reverse, ATGCATCAAACAACTGTGGCCA; *RORC*: forward, TCTGGAGCTGGCCTTTCATCATCA, reverse, TCTGCTCACTTCCAAAGAGCTGGT; and GAPDH: forward TGCACCACCAACTGCTTAG, reverse, GGATGCAGGGATGATGTTC. The reaction of PCR was performed according to the following protocol: 95 °C for 2 min, followed by 40 cycles of 95 °C for 10 sec, a specific annealing temperature for 10 sec, and 72 °C for 15 sec. Relative gene expression was calculated by 2(−Delta Delta CT) method using GAPDH as an endogenous control.

### ELISA for detecting antibody titer

For detecting anti-HBVSA antibody titer (total binding), a commercialized ELISA kit was used (Shanghai Kehua Bio-engineering Co.). In brief, plasma was diluted 10 times and incubated with HBVSA pre-coated ELISA plate for 30 min under 37°C. Then HBVSA-HRP was added for another 30 min incubation, followed by 5 washes and substrate solution was used to determine the OD450 value. The antibody titer was calculated according to the standard curve generated by the standard with a known antibody titer. For detecting anti-HBVSA antibody titer (total IgG), plasma was diluted 10 times and incubated with HBVSA pre-coated ELISA plate for 30 min under 37°C. Then the plate was washed 3 times, and added by anti-human IgG-HRP (1:60000 dilution, Sigma) for 30 min incubation under 37°C. The plate was then washed 5 times, and substrate solution was used to determine the OD450 value. IgG specific to SARS-CoV-2 spike (S) and nucleocapsid (N) proteins in plasma were measured using chemiluminescent immunoassay kits (Yhlo Biotech Co) as previously described (59). For detecting the anti-NP antibody titer, mouse serum was diluted 2,000 times and incubated with NP2-BSA or NP23-BSA pre-coated ELISA plate for 1h at RT, followed by 3 washes and incubated with anti-mouse total IgG-HRP antibody for 1h at RT. The plate was then washed 5 times, and TMB chromogen solution was used to determine the OD405 value with 0.1% SDS as the stop solution.

### *In vitro* survival and proliferation assays

For *in vitro* apoptosis assay, FACS purified CD4^+^ T cells were resuspended in complete RPMI media (10% deactivated FBS (v/v), 100 units/mL penicillin, 100 μg/mL streptomycin, 1 mM sodium pyruvate, 1% MEM nonessential amino acids (v/v), and 0.055 mM β-Mercaptoethanol in RPMI 1640 with L-glutamine and 25 mM HEPES) and cultured for 3 days, followed by Annexin V and zombie aqua (Biolegend) staining by FACS. For *in vitro* proliferation assay, FACS purified CD4^+^ T cells were labelled by 5 µM CFSE (Thermo Fisher Scientific) for 5 min, washed and seeded on 96-well U-bottom plate with T cell activation Dynabeads (Thermo Fisher Scientific) at the ratio of 3:1 (cell number:bead number) to culture for 2.5 days, followed by FACS to determine the fluorescence of CFSE. Division indices were calculated according to the online tutorial by Flowjo.

### T cell stimulation assay

To evaluate the effect of TCR stimulation on CXCR3 and CCR6 expression by Tfh cells, sorted cTfh cell subsets (2 × 10^4^/well) were stimulated with plate-bound αCD3 (5 µg/mL) and αCD28 (2 µg/mL) or rested for 18h in complete RPMI media. CXCR3 and CCR6 expression by Tfh cell subsets were analysed by FACS.

### Flow cytometry analysis

Surface staining was conducted by incubating the cells with the antibodies under room temperature for 30 min in FACS buffer (PBS + 2% FBS). For staining of intracellular cytokines, human cells were stimulated with PMA and ionomycin (500 ng/mL, eBioscience) in the presence of GolgiPlus and GolgiStop (BD Biosciences) for 4 h at 37°C. After surface staining, cells were permeabilised using Cytofix/Cytoperm (BD Biosciences). Antibodies specific to cytokines were incubated with cells for 30 min at 4°C. For intranuclear staining, surface staining was performed followed by fix/perm (eBioscience) and stained for nuclear proteins under room temperature for 45 min. Flow cytometry was performed on a FACS analyser (Fortessa X-20, BD) and the data were analyzed by FlowJo (TreeStar). The details for antibodies were shown in Table S2.

### RNA-seq data analysis

0.5-1 million naïve, T_CM_, T_EM_, Tfh_CM_, and Tfh_EM_ cells were sorted and extracted total RNA was sequenced by the Illumina platform, and the generated pair-end reads were processed online under the Galaxy project according to a standardised pipeline (60). The count files were analysed according to a published pipeline (61) for cpm normalization, MDS plot generation and differentially expressed genes calculation (low count genes were removed by filterByExpr). The heatmap was visualized by HemI. Gene set enrichment analysis (GSEA) was performed by fgsea to calculate GSEA *p* value and normalized enrichment score.

### 10x single-cell RNA-seq analysis

R script for this analysis was provided in the supplementary file. In brief, the processed Seurat object was downloaded from GSE152522 and loaded into Seurat package (62). Unsupervised clustering was then performed to extract CXCR5-expression cTfh clusters. Then Tfh1 and Tfh17 signature scores were calculated for each cell by AddModuleScore function based on the signature gene sets for Tfh1 and Tfh17 derived from GSE123812. For TCR clonality analysis, cells sharing the same TCR alpha and beta chain CDR3 amino acid sequences were assigned to the same clonotype and the clonal abundance was calculated and ranked. Finally, the Tfh1 and Tfh17 signature scores for the abundant TCR clones (abundance ≥10) were extracted for statistical analysis and visualization.

### Cell transfer and immunisation

For adoptive transfer of *in vitro* differentiated OT-II cells, CD44^+^ OT-II cells cultured under iTh0 or iTfh1/2/17 conditions were FACS-purified and 5 × 10^4^ cells were transferred into each CD28KO recipient mice, followed by OVA or NP-OVA in alum immunisations. For immunization, 50 µg ovalbumin (OVA) or NP-OVA was emulsified in alum (volume ratio 1:1) and injected through intraponeal for a dose of 200 µL per mouse. The spleens were collected on day 7 after immunisation or otherwise indicated on the paper.

### *In vitro* differentiation for OT-II cells

The method to differentiate iTfh1, iTfh2 and iTfh17-polarized cells *in vitro* was developed based on our previous paper (19). In brief, red blood cell lysed splenocytes from WT mice were left untreated (for differentiating iTh0/1/2/17) or pre-treated by 1 μg/mL lipopolysaccharide (LPS) for 24h in the complete RPMI media. 5 × 10^5^ per well LPS pre-treated splenocytes were co-cultured with FACS purified OT-II cells at the ratio of 50:1 in the presence of 1 μg/mL OVA_323-339_ peptide and indicated cytokines (Table S3) for 72h to differentiate iTfh1/2/17 cells. No cytokines were added for differentiating Th0 cells. Neutralizing antibodies anti-IL-4, anti-IFN-γ and anti-TGF-β (BioxCell) were used at 10 μg/mL. Cytokines were purchased from PeproTech.

### Statistical analysis

For human result analysis, data were not assumed Gaussian distributed thus comparisons between two groups were performed by two-tailed Wilcoxon matched-pairs signed rank test and multiple comparisons were performed by Friedman test. For mouse result analysis, data were assumed Gaussian distributed thus comparisons between two groups were performed by two-tailed unpaired *t*-test and multiple comparisons were performed by either one-way or two-way ANOVA test as specified in this paper. Corrections were not applied for multiple comparison tests because comparisons in this study were planned with specific hypotheses specified in advance. Statistical analysis was performed by Prism 9.0 software (GraphPad). *P* values < 0.05 were considered significant.

## Acknowledgements

We thank supports from Harpreet Vohra and Michael Devoy of the Imaging and Cytometry Facility at the Australian National University. The authors thank the funding supports by Australian National Health and Medical Research Council (NHMRC, GNT2009554, GNT2000466), the Bellberry-Viertel Senior Medical Research Fellowship, ANU and UQ Intramural Funding to D.Y., NHMRC grants (GNT1158404 to I.A.C., GNT1173871 to K.K., and GNT1194036 to T.H.O.N.), the National Natural Science Foundation of China grants (82130030 and 81920108011 to Z.L. and 82101198 to Y.Yao), the National Key Research and Development Program of China (2017YFC0909003) to L.Lu, and Shandong Provincial Natural Science Foundation (ZR2020ZD41, 2021ZDSYS12) to Y.W.. Part of this research was carried out at the Translational Research Institute, Woolloongabba, QLD 4102, Australia. The Translational Research Institute is supported by a grant from the Australian Government. The funders of the study had no involvement in the study design, data collection, data analysis, interpretation, writing of the report, or decision to submit the paper for publication

## Author contributions

D. Y., W.L., I.A.C. and X.G. conceived and designed the study. X.G. performed the experiments and analyzed the data with the help of Q.Z., L.X. (animal experiments), K.L., Y.Yao, T.H.O.N., L.F.A. (human data analysis), Y.Yao, D.W., Y.W., J.D., X.D., D.G., L.Lin, S.L., Y.L., L.Liu (human sample preparation), Y.Yang, K.P. (RNA-seq data analysis). K.K., Y.J., M.D., W.C., L.Lu, N.S. and Z.L. provided critical feedback on the project and manuscript, X.G. and D.Y. wrote the paper. All authors approved the final paper.

## Data and materials availability

RNA-seq dataset was accessible at GSE167309. Original R code for single-cell RNA-seq analysis can be found in the Supplementary Materials

---

**Figure S1.**
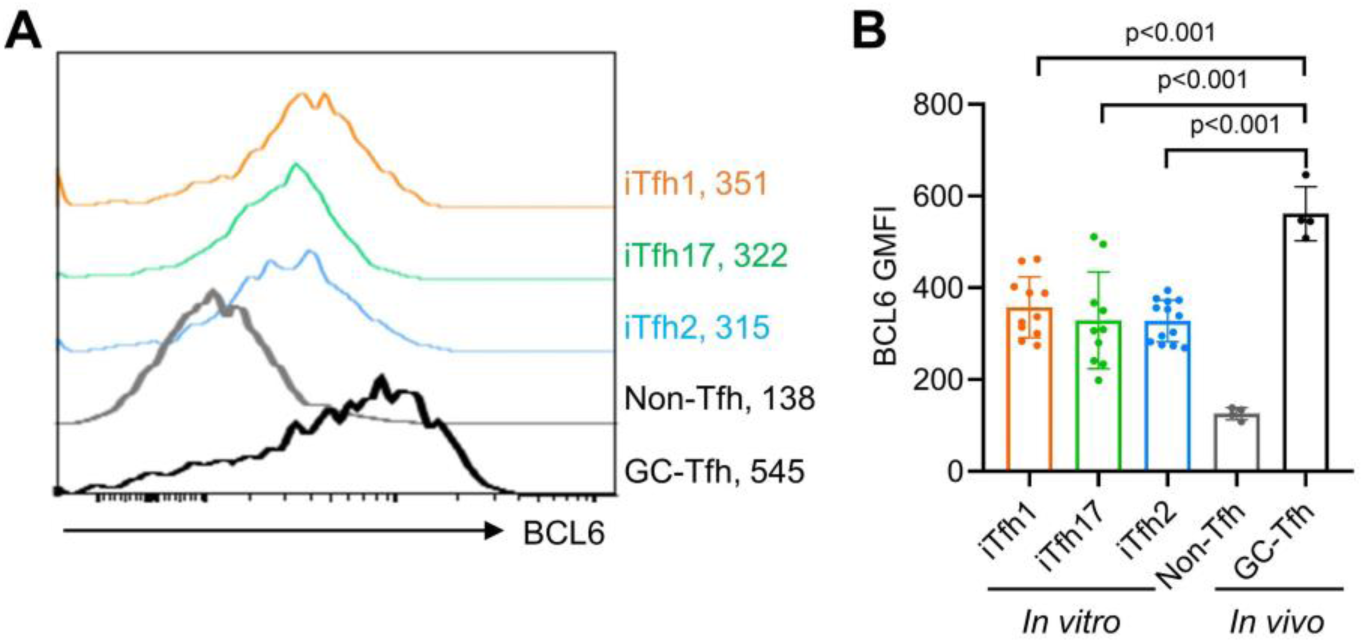
iTfh1/2/17 cells show lower BCL6 expression than GC-Tfh cells. OT-II cells were transferred into congenic WT mice followed by OVA in alum immunization. On day7, the BCL6 expressions for splenic OT-II Non-Tfh (CD44^+^CXCR5^-^PD-1^-^) and GC-Tfh cells (CD44^+^CXCR5^high^PD-1^high^) were analysed and compared with iTfh1/2/17 cells. Representative FACS plots (A) and statistics (B) comparing the expressions of BCL6 of indicated cell populations. The numbers in (A) indicate GMFI values. The results were pooled from 2 independent experiments. The *p* values were calculated by one-way ANOVA.

**Figure S2.**
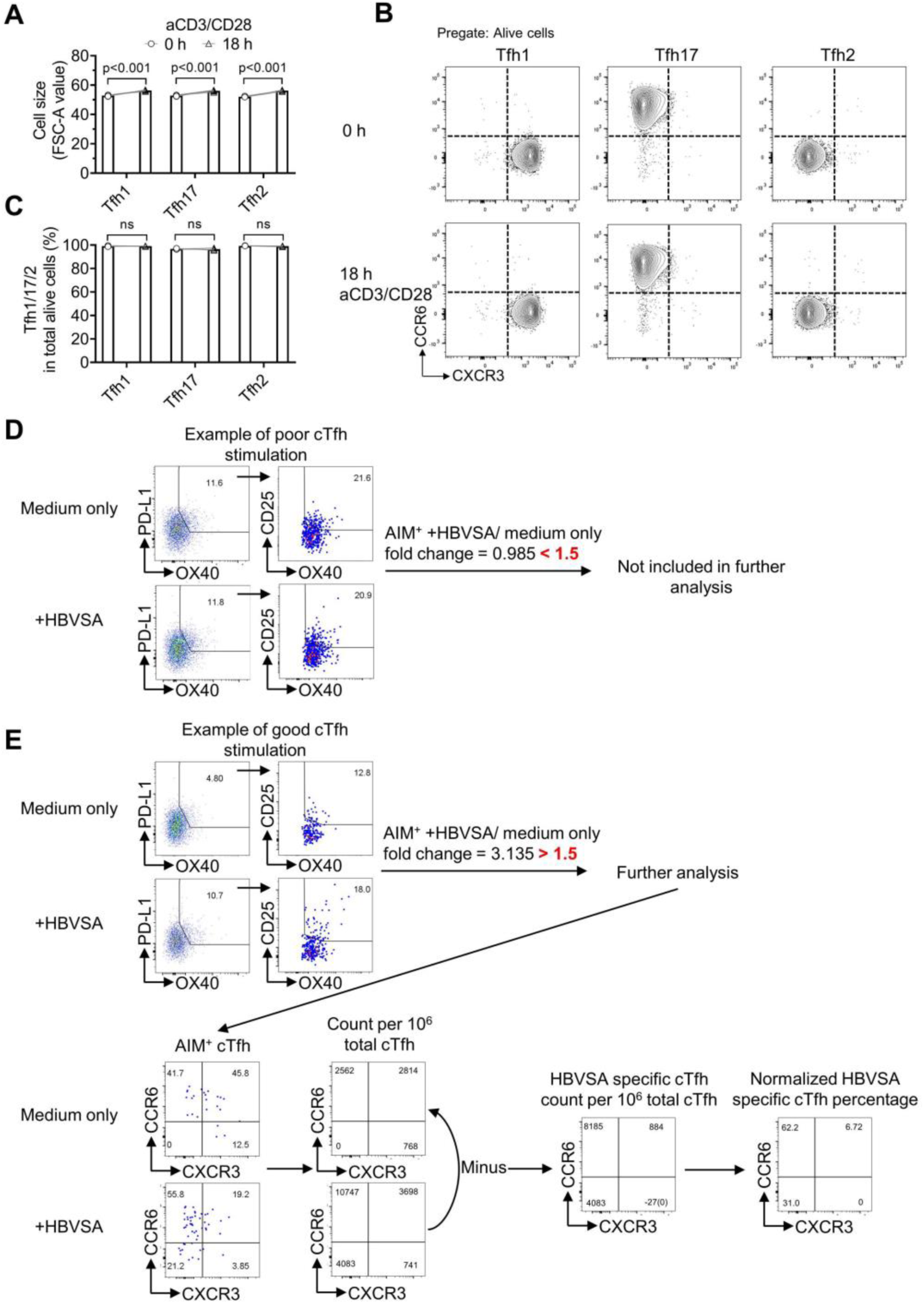
AIM assays to identify antigen-specific cTfh cells. (A-C) FACS-purified Tfh1/2/17 cells from 5 individuals were stimulated by αCD3/CD28 for 18 hours and were analysed by FACS. Statistics (A) showing the cellular sizes before/after αCD3/CD28 stimulation. Representative FACS plots (B) and statistics (C) showing the percentages of Tfh1/2/17 cells before/after stimulation. (D-E) PBMC were cultured with or without HBVSA for 18h followed by FACS analysis. Example (D) of one sample excluded from the analysis. Example (E) of one sample included in the analysis. The numbers of total cTfh cells were then normalized to 10^6^ and the absolute counts of (AIM^+^) PD-L1^+^OX40^+^CD25^+^ cTfh cells were calculated, and the absolute counts of antigen-specific Tfh1, Tfh2, Tfh17 and Tfh1/17 were obtained. Any count values less than 0 were changed to 0 because cell count cannot be negative. At last, the normalized percentage was calculated based on the absolute count of antigen-specific Tfh1, Tfh2, Tfh17 and Tfh1/17. The *p* values were calculated by paired *t*-tests.

**Figure S3.**
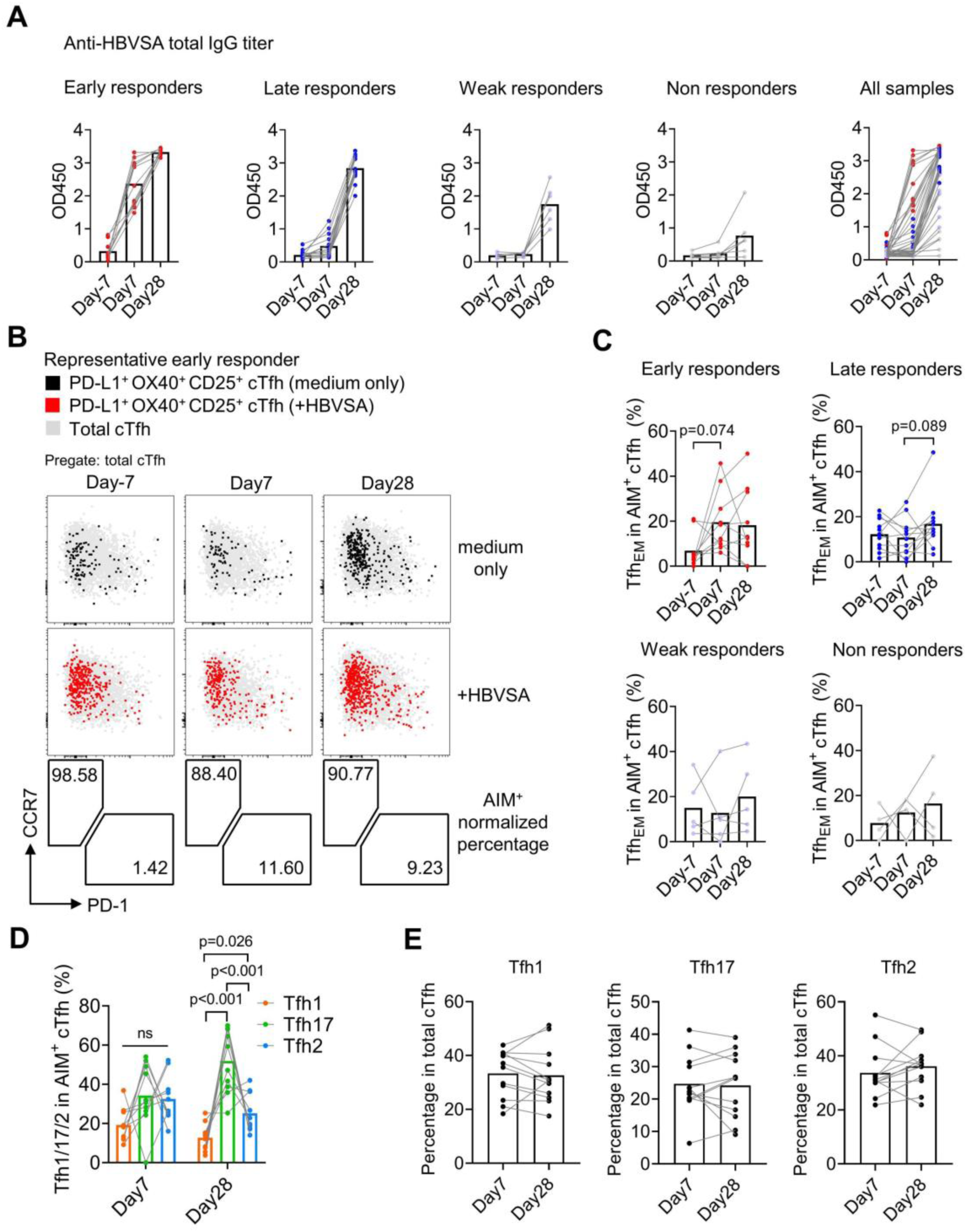
The cTfh responses in HBV vaccinated individuals. (A) Blood samples were obtained from 38 individuals who received HBV vaccines on day -7, day 7 and day 28 before/after HBV vaccination. ELISA was performed to determine the titer of anti-HBVSA IgG antibody in serum. Participants were separated into 4 groups according to their responses to the vaccine. (B-E) The PBMC were isolated and cultured for 18h with or without 20 µg/mL HBVSA, followed by FACS analysis. Representative FACS plot of one early responder (B) showing the percentage of Tfh_CM_ and Tfh_EM_ in HBVSA-specific cTfh cells on indicated time points. The statistics (C) showing the percentages of HBVSA-specific Tfh_EM_ cells on indicated time points in 4 groups of responders. The statistics of early responders (D) showing the percentages of HBVSA-specific Tfh1/2/17 cells on day 7 and day 28 post HBV vaccination. The statistics of early responders (E) showing the percentages of Tfh1/2/17 in total cTfh on day 7 and day 28 post HBV vaccination. The *p* values were calculated by Friedman test.

**Figure S4.**
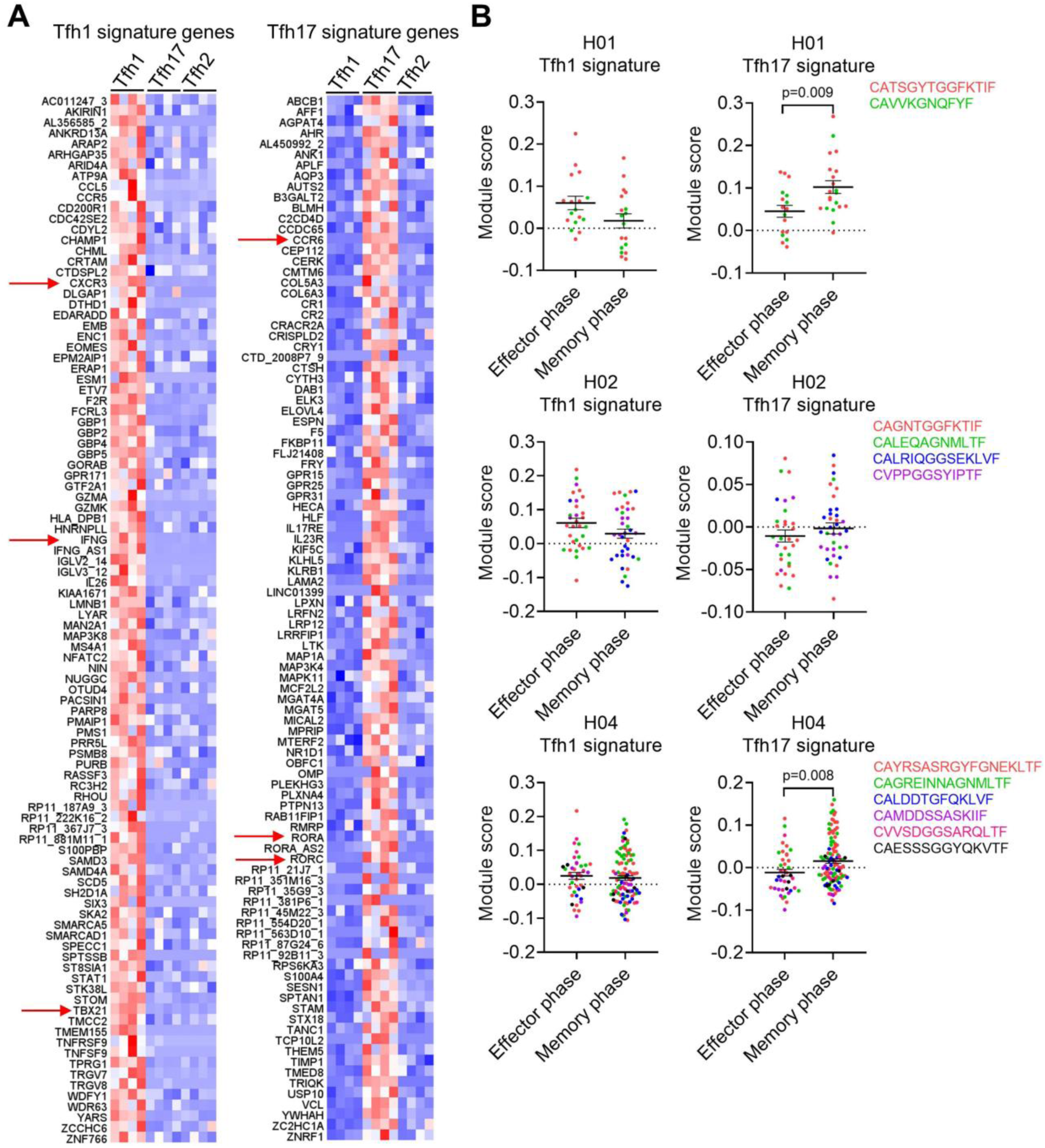
scRNA-seq analysis for influenza-specific cTfh cells. (A) Top 100 signature genes for Tfh1 and Tfh17 were generated by Limma package based on bulk RNA-seq dataset GSE123812. The heatmap showing the expressions of Tfh1 and Tfh17 signature genes in Tfh12/17 samples. (B) Tfh1 and Tfh17 signature scores in were separated by individuals (H01, H02 and H04) and each clone was colour- coded accompanied by the CDR3 amino acid sequences. The statistics comparing the Tfh1 and Tfh17 signature scores between effector and memory phase influenza-specific cTfh cells. One individual (H03) was not included since lacking abundant clones. The *p* values were calculated by student *t*-tests.

**Figure S5.**
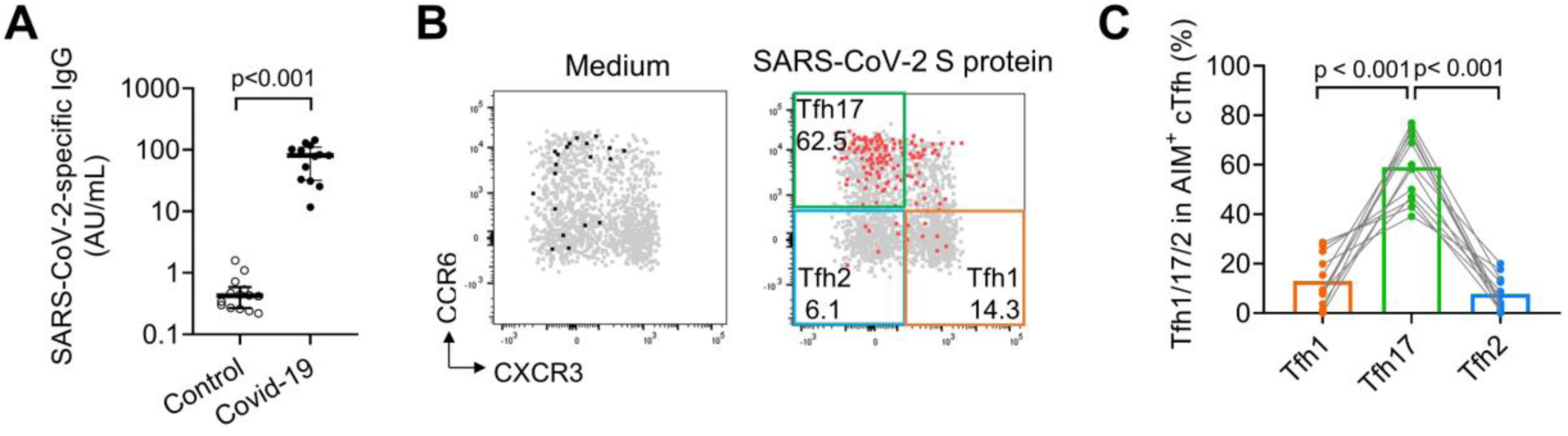
SARS-CoV-2-specific Tfh17 cells have superior persistence. Peripheral blood was collected from 14 healthy donors and 13 convalescent Covid-19 patients at 10-12 months post the infection, and ELISA or AIM assays were performed. The SARS-CoV-2 specific IgG antibody titers were shown in (A), representative FACS plots and statistics for Tfh1/2/17’s percentages in AIM^+^ cTfh in convalescent Covid-19 patients were shown in (B, C). One samples with poor signals in AIM assay were excluded. The *p* values were calculated by Friedman test.

**Table S1.** Demographics for all Human Samples Included in the Research

| Cohort description | Number | Gender<br>(female, male) | Age<br>(median, range) | Corresponding<br>Figures |
| --- | --- | --- | --- | --- |
| Healthy individuals for<br>cTfh phenotyping | 33 | 26/7 | 35 (21-71) | Figure 3A-C<br>Figure 8D-F<br>Figure S2A-C |
| Healthy individuals<br>received HBV vaccines | 38 | 8/29 | 19 (18-20) | Figure 5<br>Figure 8D-F<br>Figure S2D-E<br>Figure S3 |
| Healthy individuals for<br>measles and TT AIM<br>assay | 20 | 11/9 | 24 (18-32) | Figure 8D-F |
| Healthy children | 18 | 14/4 | 6 (0.5-12) | Figure 8D-F |
| Cord blood | 5 | 2/3 | 0 (0-0) | Figure 8D-F |
| Recovered Covid-19<br>patients | 13 | 9/4 | 33 (23-52) | Figure 6E-G |
| Healthy individuals for<br>qPCR and cytokine<br>assay | 14 | 9/5 | 42.5 (27-51) | Figure 3D-G<br>Figure 6E |

**Table S2.** Flow Cytometry Antibodies

| Antibody (human) | Clone | Antibody (mouse) | Clone |
| --- | --- | --- | --- |
| CD4 | RPA-T4 | B220 | RA3-6B2 |
| CD45RA | HI100 | CD38 | 90 |
| CXCR5 | J252D4 | CCR7 | 4B12 |
| CXCR3 | G025H7 | GL7 | GL7 |
| CCR6 | G034E3 | CD4 | RM4-4 |
| CCR7 | G043H7 | CD44 | IM7 |
| PD-1 | A17188B | CXCR5 | L138D7 |
| PD-L1 | 29E.2A3 | PD-1 | 29F.1A12 |
| OX40 | Ber-ACT35 (ACT35) | CCR7 | 4B12 |
| CD25 | BC96 | CXCR3 | CXCR3-173 |
| CD19 | HIB19 | CCR6 | 29-2L17 |
| IFN-γ | B27 | T-bet | 4B10 |
| IL-4 | MP4-25D2 | GATA3 | 16E10A23 |
| IL-17A | BL168 | RORγt | Q31-378 |
|  |  | CD45.2 | 104 |

**Table S3.** Conditions for Differentiating iTfh0, iTfh1, iTfh2 and iTfh17 Cells

| **Cell type** | **Cytokines** | **Neutralizing antibodies** |
| --- | --- | --- |
| **iTh0** | No cytokines | No antibodies |
| **iTh1** | 20ng/ml IL-12 | Anti-IL-4, anti-TGF-β |
| **iTh2** | 50ng/ml IL-4 | Anti-IFN-γ, anti-TGF-β |
| **iTh17** | 50ng/ml IL-6, 2ng/ml TGF-β | Anti-IFN-γ, anti-IL-4 |
| **iTfh1** | 100ng/ml IL-6, 50ng/ml IL-21, 1ng/ml IL-12 | Anti-IL-4, anti-TGF-β |
| **iTfh2** | 100ng/ml IL-6, 50ng/ml IL-21, 20ng/ml IL-4 | Anti-IFN-γ, anti-TGF-β |
| **iTfh17** | 100ng/ml IL-6, 50ng/ml IL-21, 0.1ng/ml TGF-β | Anti-IFN-γ, anti-IL-4 |

## References

1. Vinuesa CG, Linterman MA, Yu D, MacLennan IC. Follicular Helper T Cells. Annu Rev Immunol. 2016;34:335–68.

2. Crotty S. Follicular helper CD4 T cells (TFH). Annu Rev Immunol. 2011;29:621–63.

3. Hale JS, Youngblood B, Latner DR, Mohammed AU, Ye L, Akondy RS, et al. Distinct memory CD4+ T cells with commitment to T follicular helper- and T helper 1-cell lineages are generated after acute viral infection. Immunity. 2013;38(4):805–17.

4. He J, Tsai LM, Leong YA, Hu X, Ma CS, Chevalier N, et al. Circulating precursor CCR7(lo)PD-1(hi) CXCR5(+) CD4(+) T cells indicate Tfh cell activity and promote antibody responses upon antigen reexposure. Immunity. 2013;39(4):770–81.

5. MacLeod MK, David A, McKee AS, Crawford F, Kappler JW, Marrack P. Memory CD4 T cells that express CXCR5 provide accelerated help to B cells. J Immunol. 2011;186(5):2889–96.

6. Sage PT, Alvarez D, Godec J, von Andrian UH, Sharpe AH. Circulating T follicular regulatory and helper cells have memory-like properties. J Clin Invest. 2014;124(12):5191–204.

7. Weber JP, Fuhrmann F, Hutloff A. T-follicular helper cells survive as long-term memory cells. Eur J Immunol. 2012;42(8):1981–8.

8. Yu D, Walker LSK, Liu Z, Linterman MA, Li Z. Targeting TFH cells in human diseases and vaccination: rationale and practice. Nat Immunol. 2022.

9. Tsai LM, Yu D. Follicular helper T-cell memory: establishing new frontiers during antibody response. Immunol Cell Biol. 2014;92(1):57–63.

10. Morita R, Schmitt N, Bentebibel SE, Ranganathan R, Bourdery L, Zurawski G, et al. Human blood CXCR5(+)CD4(+) T cells are counterparts of T follicular cells and contain specific subsets that differentially support antibody secretion. Immunity. 2011;34(1):108–21.

11. Zhao J, Chen Y, Zhao Q, Shi J, Yang W, Zhu Z, et al. Increased circulating Tfh17 and PD-1+Tfh cells are associated with autoantibodies in Hashimoto’s thyroiditis. Autoimmunity. 2018;51(7):352–9.

12. Akiyama M, Suzuki K, Yamaoka K, Yasuoka H, Takeshita M, Kaneko Y, et al. Brief Report: Number of Circulating Follicular Helper 2 T Cells Correlates With IgG4 and Interleukin-4 Levels and Plasmablast Numbers in IgG4-Related Disease. Arthritis & Rheumatology. 2015;67(9):2476–81.

13. Niessl J, Baxter AE, Morou A, Brunet-Ratnasingham E, Sannier G, Gendron-Lepage G, et al. Persistent expansion and Th1-like skewing of HIV-specific circulating T follicular helper cells during antiretroviral therapy. EBioMedicine. 2020;54:102727.

14. Obeng-Adjei N, Portugal S, Tran TM, Yazew TB, Skinner J, Li S, et al. Circulating Th1-Cell-type Tfh Cells that Exhibit Impaired B Cell Help Are Preferentially Activated during Acute Malaria in Children. Cell Rep. 2015;13(2):425–39.

15. Bentebibel SE, Lopez S, Obermoser G, Schmitt N, Mueller C, Harrod C, et al. Induction of ICOS+CXCR3+CXCR5+ TH cells correlates with antibody responses to influenza vaccination. Sci Transl Med. 2013;5(176):176ra32.

16. Spensieri F, Borgogni E, Zedda L, Bardelli M, Buricchi F, Volpini G, et al. Human circulating influenza- CD4+ ICOS1+IL-21+ T cells expand after vaccination, exert helper function, and predict antibody responses. Proc Natl Acad Sci U S A. 2013;110(35):14330–5.

17. Herati RS, Muselman A, Vella L, Bengsch B, Parkhouse K, Del Alcazar D, et al. Successive annual influenza vaccination induces a recurrent oligoclonotypic memory response in circulating T follicular helper cells. Sci Immunol. 2017;2(8).

18. Sallusto F, Geginat J, Lanzavecchia A. Central memory and effector memory T cell subsets: function, generation, and maintenance. Annu Rev Immunol. 2004;22:745–63.

19. Gao X, Wang H, Chen Z, Zhou P, Yu D. An optimized method to differentiate mouse follicular helper T cells in vitro. Cell Mol Immunol. 2020;17(7):779–81.

20. Read KA, Powell MD, Sreekumar BK, Oestreich KJ. In Vitro Differentiation of Effector CD4(+) T Helper Cell Subsets. Methods Mol Biol. 2019;1960:75–84.

21. Lu KT, Kanno Y, Cannons JL, Handon R, Bible P, Elkahloun AG, et al. Functional and epigenetic studies reveal multistep differentiation and plasticity of in vitro-generated and in vivo-derived follicular T helper cells. Immunity. 2011;35(4):622–32.

22. Nurieva RI, Chung Y, Hwang D, Yang XO, Kang HS, Ma L, et al. Generation of T follicular helper cells is mediated by interleukin-21 but independent of T helper 1, 2, or 17 cell lineages. Immunity. 2008;29(1):138–49.

23. Gaya M, Barral P, Burbage M, Aggarwal S, Montaner B, Warren Navia A, et al. Initiation of Antiviral B Cell Immunity Relies on Innate Signals from Spatially Positioned NKT Cells. Cell. 2018;172(3):517–33 e20.

24. Bouneaud C, Garcia Z, Kourilsky P, Pannetier C. Lineage relationships, homeostasis, and recall capacities of central- and effector-memory CD8 T cells in vivo. J Exp Med. 2005;201(4):579–90.

25. Chevalier N, Jarrossay D, Ho E, Avery DT, Ma CS, Yu D, et al. CXCR5 expressing human central memory CD4 T cells and their relevance for humoral immune responses. The Journal of Immunology. 2011;186(10):5556–68.

26. Omilusik KD, Best JA, Yu B, Goossens S, Weidemann A, Nguyen JV, et al. Transcriptional repressor ZEB2 promotes terminal differentiation of CD8+ effector and memory T cell populations during infection. J Exp Med. 2015;212(12):2027–39.

27. Bruce MG, Bruden D, Hurlburt D, Zanis C, Thompson G, Rea L, et al. Antibody Levels and Protection After Hepatitis B Vaccine: Results of a 30-Year Follow-up Study and Response to a Booster Dose. J Infect Dis. 2016;214(1):16–22.

28. Dan JM, Lindestam Arlehamn CS, Weiskopf D, da Silva Antunes R, Havenar-Daughton C, Reiss SM, et al. A Cytokine-Independent Approach To Identify Antigen-Specific Human Germinal Center T Follicular Helper Cells and Rare Antigen-Specific CD4+ T Cells in Blood. J Immunol. 2016;197(3):983–93.

29. Reiss S, Baxter AE, Cirelli KM, Dan JM, Morou A, Daigneault A, et al. Comparative analysis of activation induced marker (AIM) assays for sensitive identification of antigen-specific CD4 T cells. PLoS One. 2017;12(10):e0186998.

30. Meckiff BJ, Ramirez-Suastegui C, Fajardo V, Chee SJ, Kusnadi A, Simon H, et al. Imbalance of Regulatory and Cytotoxic SARS-CoV-2-Reactive CD4(+) T Cells in COVID-19. Cell. 2020;183(5):1340–53 e16.

31. Yost KE, Satpathy AT, Wells DK, Qi Y, Wang C, Kageyama R, et al. Clonal replacement of tumor-specific T cells following PD-1 blockade. Nat Med. 2019;25(8):1251–9.

32. Huang J, Ruan S, Wu X, Zhou X. Seasonal transmission dynamics of measles in China. Theory Biosci. 2018;137(2):185–95.

33. Van Damme P, Dionne M, Leroux-Roels G, Van Der Meeren O, Di Paolo E, Salaun B, et al. Persistence of HBsAg-specific antibodies and immune memory two to three decades after hepatitis B vaccination in adults. J Viral Hepat. 2019;26(9):1066–75.

34. Nanan R, Rauch A, Kämpgen E, Niewiesk S, Kreth HW. A novel sensitive approach for frequency analysis of measles virus-specific memory T-lymphocytes in healthy adults with a childhood history of natural measles. Journal of General Virology. 2000;81(5):1313–9.

35. Locci M, Havenar-Daughton C, Landais E, Wu J, Kroenke MA, Arlehamn CL, et al. Human circulating PD-1+CXCR3-CXCR5+ memory Tfh cells are highly functional and correlate with broadly neutralizing HIV antibody responses. Immunity. 2013;39(4):758–69.

36. Deng J, Wei Y, Fonseca VR, Graca L, Yu D. T follicular helper cells and T follicular regulatory cells in rheumatic diseases. Nat Rev Rheumatol. 2019;15(8):475–90.

37. Yao Y, Chen CL, Yu D, Liu Z. Roles of follicular helper and regulatory T cells in allergic diseases and allergen immunotherapy. Allergy. 2021;76(2):456–70.

38. Baiyegunhi O, Ndlovu B, Ogunshola F, Ismail N, Walker BD, Ndung’u T, et al. Frequencies of Circulating Th1-Biased T Follicular Helper Cells in Acute HIV-1 Infection Correlate with the Development of HIV-Specific Antibody Responses and Lower Set Point Viral Load. J Virol. 2018;92(15).

39. Rydyznski Moderbacher C, Ramirez SI, Dan JM, Grifoni A, Hastie KM, Weiskopf D, et al. Antigen-Specific Adaptive Immunity to SARS-CoV-2 in Acute COVID-19 and Associations with Age and Disease Severity. Cell. 2020;183(4):996–1012 e19.

40. Dan JM, Mateus J, Kato Y, Hastie KM, Yu ED, Faliti CE, et al. Immunological memory to SARS-CoV-2 assessed for up to 8 months after infection. Science. 2021;371(6529).

41. Gowthaman U, Chen JS, Zhang B, Flynn WF, Lu Y, Song W, et al. Identification of a T follicular helper cell subset that drives anaphylactic IgE. Science. 2019;365(6456).

42. Hirota K, Turner JE, Villa M, Duarte JH, Demengeot J, Steinmetz OM, et al. Plasticity of Th17 cells in Peyer’s patches is responsible for the induction of T cell-dependent IgA responses. Nat Immunol. 2013;14(4):372–9.

43. McCarron MJ, Marie JC. TGF-beta prevents T follicular helper cell accumulation and B cell autoreactivity. J Clin Invest. 2014;124(10):4375–86.

44. Marshall HD, Ray JP, Laidlaw BJ, Zhang N, Gawande D, Staron MM, et al. The transforming growth factor beta signaling pathway is critical for the formation of CD4 T follicular helper cells and isotype-switched antibody responses in the lung mucosa. Elife. 2015;4:e04851.

45. Young B, Sadarangani S, Jiang L, Wilder-Smith A, Chen MI. Duration of Influenza Vaccine Effectiveness: A Systematic Review, Meta-analysis, and Meta-regression of Test-Negative Design Case-Control Studies. J Infect Dis. 2018;217(5):731–41.

46. Dan JM, Mateus J, Kato Y, Hastie KM, Yu ED, Faliti CE, et al. Immunological memory to SARS-CoV-2 assessed for up to 8 months after infection. Science. 2021:eabf4063.

47. Chan J-A, Loughland JR, de Labastida Rivera F, SheelaNair A, Andrew DW, Dooley NL, et al. Th2-like T Follicular Helper Cells Promote Functional Antibody Production during Plasmodium falciparum Infection. Cell Reports Medicine. 2020;1(9).

48. Goel RR, Painter MM, Apostolidis SA, Mathew D, Meng W, Rosenfeld AM, et al. mRNA vaccines induce durable immune memory to SARS-CoV-2 and variants of concern. Science. 2021;374(6572):abm0829.

49. Wragg KM, Lee WS, Koutsakos M, Tan H-X, Amarasena T, Reynaldi A, et al. Establishment and recall of SARS-CoV-2 spike epitope-specific CD4+ T cell memory. Nature Immunology. 2022;23(5):768–80.

50. Tauzin A, Nayrac M, Benlarbi M, Gong SY, Gasser R, Beaudoin-Bussières G, et al. A single dose of the SARS-CoV-2 vaccine BNT162b2 elicits Fc-mediated antibody effector functions and T cell responses. Cell host & microbe. 2021;29(7):1137–50. e6.

51. Lindenstrom T, Woodworth J, Dietrich J, Aagaard C, Andersen P, Agger EM. Vaccine-induced th17 cells are maintained long-term postvaccination as a distinct and phenotypically stable memory subset. Infect Immun. 2012;80(10):3533–44.

52. Kryczek I, Zhao E, Liu Y, Wang Y, Vatan L, Szeliga W, et al. Human TH17 cells are long-lived effector memory cells. Sci Transl Med. 2011;3(104):104ra0.

53. Muranski P, Borman ZA, Kerkar SP, Klebanoff CA, Ji Y, Sanchez-Perez L, et al. Th17 cells are long lived and retain a stem cell-like molecular signature. Immunity. 2011;35(6):972–85.

54. Sun Z, Unutmaz D, Zou Y-R, Sunshine MJ, Pierani A, Brenner-Morton S, et al. Requirement for RORγ in Thymocyte Survival and Lymphoid Organ Development. Science. 2000;288(5475):2369.

55. Yao Y, Chen Z, Zhang H, Chen C, Zeng M, Yunis J, et al. Selenium-GPX4 axis protects follicular helper T cells from ferroptosis. Nat Immunol. 2021;22(9):1127–39.

56. Chen Z, Wang N, Yao Y, Yu D. Context-dependent regulation of follicular helper T cell survival. Trends Immunol. 2022;43(4):309–21.

57. Sahin U, Muik A, Derhovanessian E, Vogler I, Kranz LM, Vormehr M, et al. COVID-19 vaccine BNT162b1 elicits human antibody and TH1 T cell responses. Nature. 2020;586(7830):594–9.

58. Lederer K, Castano D, Gomez Atria D, Oguin TH, 3rd, Wang S, Manzoni TB, et al. SARS-CoV-2 mRNA Vaccines Foster Potent Antigen-Specific Germinal Center Responses Associated with Neutralizing Antibody Generation. Immunity. 2020;53(6):1281–95 e5.

59. Yao Y, Wang ZZ, Huang A, Liu Y, Wang N, Wang ZC, et al. TFH2 cells associate with enhanced humoral immunity to SARS-CoV-2 inactivated vaccine in patients with allergic rhinitis. Clinical and translational medicine. 2022;12(1):e717.

60. Jalili V, Afgan E, Gu Q, Clements D, Blankenberg D, Goecks J, et al. The Galaxy platform for accessible, reproducible and collaborative biomedical analyses: 2020 update. Nucleic acids research. 2020;48(W1):W395–W402.

61. Law CW, Alhamdoosh M, Su S, Dong X, Tian L, Smyth GK, et al. RNA-seq analysis is easy as 1-2-3 with limma, Glimma and edgeR. F1000Res. 2016;5.

62. Hao Y, Hao S, Andersen-Nissen E, Mauck WM, 3rd, Zheng S, Butler A, et al. Integrated analysis of multimodal single-cell data. Cell. 2021;184(13):3573–87 e29.

